# Dosage compensation defects due to *roX* RNA deletion are rescued by recalibration of X/autosome stoichiometry

**DOI:** 10.64898/2026.02.24.707606

**Authors:** Fotios Gkountromichos, George Yankson, Muhunden Jayakrishnan, Aline Campos Sparr, Marisa Müller, Patrick Heun, Peter B. Becker

**Author notes:** To whom correspondence should be addressed: Peter B. Becker.

## Abstract

Metazoa evolved regulatory networks to balance the expression of their sex chromosomes. In *Drosophila*, males have a single gene-rich X chromosome, whereas females have two. Balanced X/autosome expression is essential for viability, and in male flies is achieved by activation of genes on the X through the male-specific-lethal (MSL) dosage compensation complex (DCC). This ribonucleoprotein assembly contains long, non-coding *roX* RNAs.

To dissect the functional requirements of *roX* in a cell-based system, we deleted the *roX2* gene in male S2 cells and selected two independent lines lacking detectable *roX* RNA. In the absence of *roX*, the remaining MSL protein complex was unable to associate with known or newly identified binding sites and thus failed to activate transcription. Surprisingly, the X/autosome expression ratio appeared nevertheless compensated. Cytogenetic and genomic analyses revealed that both *roX*-deficient cell populations had acquired additional X chromosomes. Apparently, chromosome gains due to mis-segregation made up for the loss of DCC-mediated dosage compensation. Interestingly, ectopic expression of full-length *roX2*, but not of shortened derivatives, fully restored DCC binding and normalized the karyotype. These findings illustrate that X chromosome dosage compensation is critical for viability even in cultured cells and provide a striking example of rapid evolution under stringent selection.

**Graphical abstract:** 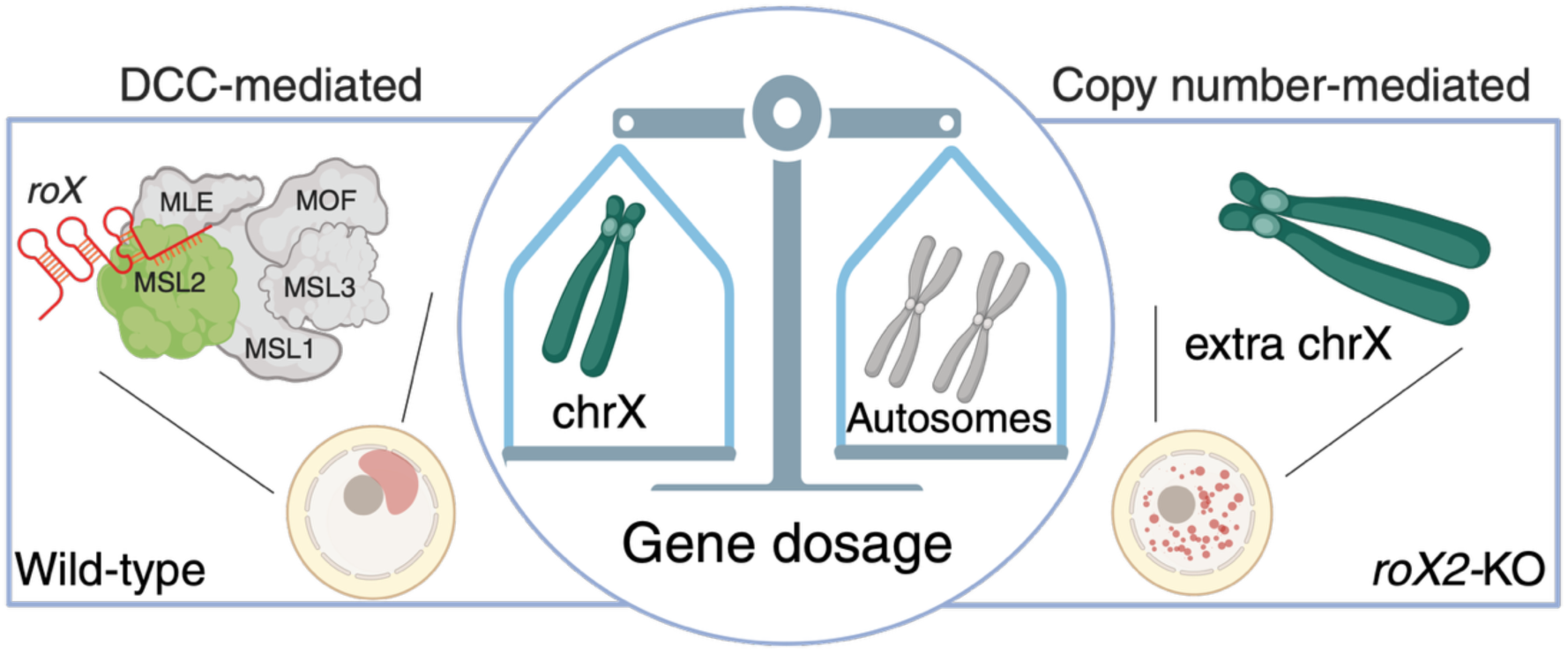

## Introduction

Maintaining balanced gene expression between sex chromosomes and autosomes is of vital importance. In *Drosophila melanogaster* the proteins encoded by male-specific-lethal (MSL) genes form a ribonucleoprotein complex (the dosage compensation complex, DCC) that transcriptionally compensates the X chromosome monosomy in males flies. An epigenetic reader/writer module composed of MSL3 and MOF is connected via the scaffold protein MSL1 to the DNA binding subunit, MSL2, which assures the selective targeting of the DCC to the X chromosome. MSL3 binds to actively transcribed chromatin marked by methylated histone H3 at lysine 36 (H3K36me). The acetyltransferase MOF then acetylates histone H4 lysine 16 (H4K16ac), which prevents chromatin compaction. This activity contributes to an approximate doubling of transcription from the male X chromosome, thus matching the combined output of the two female X chromosomes (1,2). In the absence of a functional DCC-mediated dosage compensation system, male *Drosophila* embryos die during gastrulation (3). Whether this is caused by haploinsufficient genes that are critical for further development or because of a generally compromised fitness due to unbalanced genome-wide expression, is not known.

The functions of the protein components of the DCC are, in principle, well-characterized, but the role of the long, non-coding *roX* (RNA-on-X) RNAs that associate with the MSL complex *in vivo* remains poorly understood. The realization that the long *roX1* and the much shorter *roX2* RNAs tightly colocalized with the MSL proteins on the X chromosome suggested an involvement in dosage compensation, which was then confirmed (4–8). The *roX1* and *roX2* genes are located on the X chromosome. The discovery of X-linked non-coding RNAs involved in dosage compensation in *Drosophila* resonated with the critical involvement of *Xist* RNA in mammalian X inactivation, suggesting a common principle underlying diverse dosage compensation strategies (9). In contrast to *Xist*, *Drosophila roX* RNAs can trigger X chromosome activation even if transcribed ectopically from an autosome (5,10). *RoX* RNA genes evolved fast in *Drosophilids*, suggesting that they may have driven the evolution of the dosage compensation system as we know it today (10,11).

The two *roX* genes serve distinct functions during development, but each RNA alone can sustain male life, if put to the test. *RoX1* is transcribed in both sexes up to stage 12, from which point onwards it is male-specific. The male-specific transcription of *roX2* RNA begins at stage 10 and its expression mirrors the profile of gradual X chromosome activation (6,12). Deletion of either *roX* gene does not compromise male viability, but impairment of both causes male lethality, with 5% of males emerging as developmentally delayed escapers (13,14). Strangely, in these organismic studies, male larvae lacking both *roX* genes appear healthy, presumably because of maternal contribution. Overexpression of MSL proteins can partially overcome a *roX* deficiency, pointing to a role for *roX* RNA in targeting the DCC to the X chromosome, rather than being required for activation of its HAT activity (10,15). The association of either *roX* RNA with the MSL proteins is mediated by the DexH helicase MLE (maleless) (16). The helicase unwinds conserved stem-loop structures at the 3’ end of *roX* RNAs, exposing linear motifs to which MSL2 can bind (17–20). This interaction has been proposed to “trap” the DCC specifically on the X chromosome, potentially explaining the exclusive X chromosome binding despite genome-wide distribution of the DNA sequences that resemble MSL recognition elements (MREs) (10,21,22).

Currently, the highly selective association of the DCC with the X is explained by two non-exclusive models. The first model assumes that the DCC associates with the X chromosome in a 2-step process. Accordingly, the initial recognition of a limited number of DNA elements exclusive to the X, such as High Affinity sites (HAS), Chromosome Entry Sites (CES) and Pioneering on the X (PionX) sites is independent of RNA. From these sites the complex ‘spreads’ to neighboring genes in a *roX*-dependent manner (1,2). An alternative model poses that the X chromosome evolved numerous DCC binding sites with varying affinities, where higher-affinity sites are less dependent on *roX* RNA for DCC recruitment (12,23–25). Traditional methods such as polytene chromosome immunostaining and standard chromatin immunoprecipitation sequencing (ChIP-seq) primarily detect only the highest occupancy sites and cannot easily quantify *roX*-dependency at sites of varying affinity, making it difficult to distinguish between the models.

The complex developmental phenotypes of *roX* gene mutations motivated us to explore the role of *roX* RNA in a simple proliferative cell culture model. Immortalized hematopoietic *Drosophila* S2 cells (26) previously served as a useful tool to study dosage compensation mechanisms (10,27–29). Despite significant aneuploidy, S2 cells compensate for the reduced dose of the X chromosome (relative to autosomes) by two additive mechanisms: approximately half of the required transcription boost is achieved through a generic, mechanistically unclear, feedback response to chromosomal imbalance, while the remaining activation depends on the DCC (28). Curiously, RNAi-mediated depletion of MSL proteins disrupts localization of the DCC but does not impair cell proliferation over the course of the experiment, typically 6-10 days (27–29). It is possible that dosage compensation is critical for *Drosophila* development but dispensable for proliferation of these cells in culture.

We used CRISPR-Cas9 genome editing to generate S2 cell lines lacking all *roX* expression, in order to investigate the structural and functional requirements of this long non-coding RNA for dosage compensation. In the absence of *roX* RNAs, the MSL proteins form a complex yet fail to localize to the X chromosome (20). The *roX*-deficient single-cell model enables quantitative evaluation of *roX* function through phenotypic rescue. While full-length *roX2* complements all deficiencies, shorter derivatives do not, establishing a tractable system for comprehensive structure-function dissection of this highly evolved lncRNA.

Using CUT&RUN with improved informatic csaw-based analysis we expanded the MSL2 binding site catalog to over 1,800 sites the vast majority X-chromosomal, including most known MREs and many novel sites, half of which are promoter-proximal. Contrary to the two-step model, which posits *roX*-independent DCC binding at PionX and HAS sites, we found that *roX* is required for DCC occupancy at all site classes. This supports a quantitative affinity hierarchy model, in which DCC spreading is achieved by successive filling of progressively lower-affinity sites rather than through qualitatively distinct classes of sites that differ in their *roX*-dependency.

Surprisingly, X-chromosomal gene expression in these *roX*-deficient cells matched wild-type levels. We discovered that up to 50% of the two independent knockout cell populations had gained extra X chromosomes, effectively compensating for the loss of DCC function through karyotypic adjustment. Remarkably, stable expression of a *roX* transgene led to loss of the extra X, strongly suggesting that dosage compensation is vital even for aneuploid cells in culture and demonstrating rapid karyotypic evolution under stringent selection.

## Materials and Methods

### Cell Culture and manipulations

*Drosophila melanogaster* S2 cells were obtained from Stefan Ameres (IMBA, Vienna). They were cultured at 26 °C in Schneider’s medium (Thermo-Fisher, Cat. No. 21720024) supplemented with 10% FBS (PAN Biotech, Cat. No. P30-3031 or Capricon Scientific, Cat. No. FBS-12A) and 1% Penicillin-Streptomycin (Sigma-Aldrich, Cat. No. P-4333). Cells were routinely counted with the CASY cell counter (Roche Innovatis).

### CRISPR-Cas9 genome editing

Genome editing was done using the CRISPR-Cas9 technology as described in (30) with some alterations. Candidate gRNAs targeting the promoter region of *roX2* were designed using Benchling (31) while for the 3’ end targeting of *roX2* sgRNAs were initially designed using the GPP sgRNA designer (https://portals.broadinstitute.org/gpp/public/analysis-tools/sgrna-design; (32)). All gRNAs were tested for off-target effects using the CRIPSRfly “Optimal target finder” (http://targetfinder.flycrispr.neuro.brown.edu, guide length = 20, Stringency = high, PAM = NGG only) (33). The 20 bp gRNAs were fused to a tracrRNA backbone during synthesis and cloned downstream of the *Drosophila* U6 promoter. These constructs were synthesized as gBlocks (Integrated DNA Technologies) and PCR-amplified before transfection using the Phusion polymerase (NEB, Cat. No. M0530L). Transfection of cells for genome editing was done with XtremeGENE HP transfection reagent (Sigma-Aldrich, Cat. No. 6366244001). S2 cells (10^6^) were seeded in 500 μl medium per well of a 24-well plate and allowed to attach for 3–4 hr. Cells were transfected with a mixture containing 190 ng of a plasmid expressing Cas9 and blasticidin resistance (pIB_Cas9_Blast)(30) and 110 ng total sgRNA plasmids (36.6 ng each of sgRNA1, sgRNA3, and sgRNA5) in 100 μl serum-free medium. XtremeGENE HP transfection reagent (1.5 μl) was added to the DNA mixture, vortexed briefly, and incubated for 35 minutes at room temperature (RT) before dropwise addition to cells. After 24 hours, medium was replaced with selection medium containing blasticidin (final concentration 25 μg/ml, Gibco, Cat. No. A1113903). After 3 days in selection medium, cells were taken out of selection and polyclonal pools were screened by PCR for the *roX2* deletion. Genomic DNA was isolated using a PCR clean-up kit (Macherey-Nagel, REF 740609.50) and PCR amplification was performed with Phusion polymerase (Thermo, Cat. No. F-530XL) using the following conditions: 98°C for 1 min; 35 cycles of 98°C for 20 sec, 65°C for 30 sec, 72°C for 1.5 min; final extension at 72°C for 10 min. PCR products were analyzed on 2% agarose gels. For more information on the sequences and primers used, see Supplementary Table 1. For clonal isolation, edited polyclonal pools were seeded at low density of 1000, 2000 or 3000 cells per 6 cm plate and allowed to attach for at least 1 hr. Cells were overlaid with a 1:1 mixture of freshly prepared 2× Schneider’s medium (Serva, Cat. No. 47521.04) and 0.8% low melting agarose (BioRad, Cat. No. 161-3111) equilibrated to 37°C (3.5 ml per plate). Plates were sealed with parafilm, placed in a humidified chamber, and incubated at 26°C for 2–3 weeks to allow clonal outgrowth. Individual colonies were picked, expanded for approximately 2 weeks, and genotyped by PCR with sequencing confirmation of deletion junctions.

### Stable transgenic cell line generation

Stable transgenic lines expressing *roX* variants were generated by transfection of *roX2* knock out S2 cells (KO-B) using Effectene transfection reagent (Qiagen, Cat. No. 301425) according to the manufacturer. Briefly, 8 x 10^5^ cells were seeded in 350 μl medium per well of a 24-well plate and allowed to adhere for 3–4 hours. Transfection complexes were prepared by mixing 200 ng of *roX* expression plasmid (*roX* being expressed under the HSP70 promoter) and 10 ng of blasticidin resistance plasmid in 60 μl Buffer EC, followed by addition of 1.6 μl Enhancer, vortexing, and incubation at RT for 2–5 min. Effectene reagent (5 μl) was then added, vortexed for 10 sec, and incubated at RT for 5–10 min. The transfection mixture was diluted with 350 μl medium and added dropwise to cells. After 3 days, medium was replaced with selection medium containing 10 μg/ml blasticidin (Gibco, Cat. No. A1113903). Cells were maintained under blasticidin selection and monitored for transgene expression for 3–4 weeks, then taken out of selection and passaged several times before use in experiments.

### Biochemical methods

#### Immunoblotting

Cells (1.5 x 10^6^) were harvested, washed once with PBS, and resuspended in 60 μl Laemmli sample buffer. Extracts were boiled at 95°C for 10 min and stored at −20°C until use. Proteins were separated by SDS-PAGE on either 8% acrylamide gels (Serva, Cat. Nos. 43260.01, 43262.01) for larger proteins or 14% gels for histones and histone marks, loading approximately 2.5 x 10^5^ cell equivalents per lane. Electrophoresis was carried out in Running Buffer (25 mM Tris, 192 mM glycine, 0.1% SDS) and proteins were transferred to nitrocellulose membranes (Cytiva, Cat. No. GE10600002) by wet transfer at 350 mA on ice for approximately 90 min. Transfer Buffer (25 mM Tris, 192 mM glycine) was supplemented with 20% methanol. Membranes were blocked in 5% BSA in TBS for 1 hr at RT, then incubated with primary antibodies diluted in TBS overnight at 4°C. Primary antibodies were used to blot for MSL2 (guinea pig α-MSL2), MOF, MLE, H4K16ac, Lamin and histone H3. Following three 10-min washes in TBS-T (0.1% Tween-20, Sigma, Cat. No. P9416), membranes were incubated with LI-COR IRDye secondary antibodies in TBS for 1 hr at RT. For a detailed list of antibodies and their dilutions used see Supplementary Table 1. After three additional washes in TBS-T, signals were detected using the LI-COR Odyssey CLx Imaging System. Band intensities were quantified using the Image Studio Lite Software (Version 5.2.5, LI-COR Biosciences) and normalized to Lamin (for non-histone proteins) or H3 (for histone modifications) and compared to WT levels. Western blot protein ratios were quantified from biological replicates (cells harvested on different days), with technical replicates representing independent gel images of each biological sample. The data were processed using R (v4.5.1)(34) For normalization, technical replicates were normalized to the mean wild-type (WT) value within the same gel image, then averaged to yield a single fold-change value per biological replicate. Outliers among biological replicates were identified using the Median Absolute Deviation (MAD) method and excluded if at least three replicates remained per genotype. Statistical comparisons between genotypes and WT were performed using two-sided t-tests on log2-transformed fold-change values of biological replicates. Significance levels: **p < 0.001, *p < 0.01, p < 0.05, ns = not significant. Plots display individual biological replicates (larger points), technical replicates (smaller faint points), means, and standard deviation error bars.

#### RNA isolation, cDNA synthesis, and Real-time quantitative PCR (RT-qPCR)

Total RNA from 4.5 x 10^6^ cells was isolated using the RNeasy kit (Qiagen, Cat. No. 74104) according to the manufacturer’s instructions. RNA concentrations were measured, and samples were stored at −80°C. For DNase treatment, 2 μg RNA was incubated with 1 μl recombinant DNase (Roche, Ref. 04716728001) in 1x DNase buffer with 1 μl of recombinant ribonuclease inhibitors (RNasin, Promega, Cat. No. N2511) in a final volume of 20 μl for 1 hr at 37°C. cDNA from 1 μg of DNase-treated RNA was synthesized using SuperScript III reverse transcriptase kit and random hexamers (Invitrogen, Cat. No. 18080051). The cDNA samples were treated with 0.5 μl RNase H (NEB, Cat. No. M0297L) at 37°C for 20 min, and finally cDNA was stored at −20°C. Quantitative PCR was performed in 384-well plates using Fast SYBR Green Master Mix (Thermo, Cat. No. 4385610) on a LightCycler 480 instrument (Roche). Each reaction contained 1x SYBR Green Master Mix, 1 μM final concentration for each of forward and reverse primers and 1:10 diluted cDNA for all primers used. For a detailed list with oligos used in this study see Supplementary Table 1. Reactions were performed in technical triplicates, with biological replicates representing cells harvested and processed on different days. Data were analyzed in R (v4.5.1) using the ΔΔCp method with GAPDH as the reference gene after averaging the technical replicates, and relative expression was calculated as 2^(−ΔΔCp), and expressed as the mean fold change over WT of n=4 biological replicates. Error bars represent the standard deviation of the biological replicates.

#### CUT&RUN and Chromatin immunoprecipitation after MNase treatment (ChIP-MNase) sequencing

CUT&RUN was performed as described previously (25,35). For normalizing MSL2, pre-immune serum (PPI) was used as a negative control. For H4K16ac, rabbit IgG was used as negative control for normalization. ChIP-seq on MNase-digested chromatin and sonicated chromatin was performed as previously described in (36) and (37). Both ChIP-MNase and CUT&RUN yielded DNA was quantified using the Qubit dsDNA HS Assay Kit (Invitrogen, Cat. No. Q32851). Libraries were prepared with NEBNext Ultra II DNA Library Prep Kit for Illumina (NEB, E7645) according to the manufacturer for ChIP and with subtle modifications for CUT&RUN (https://doi.org/10.17504/protocols.io.bagaibse, (38)). Quality control and analysis of all libraries was done with the TapeStation system (Agilent). Libraries were sequenced on NextSeq1000 (Illumina) instrument yielding typically 20–25 million for ChIP-MNase or ∼5 million for CUT&RUN paired-end reads per sample. All sequencing was performed at the Laboratory of Functional Genomic Analysis (LAFUGA, Gene Center Munich, LMU).

#### Microscopy

*Immunofluorescence microscopy* (IFM) analysis of S2 cells was performed as described in (10,37) with slight alterations. Briefly, coverslips were placed in 12-well plates and UV-sterilized for 10 min. After sterilization coverslips were coated with poly-L-lysine (Sigma #P8920, 0.01% w/v final concentration) and air-dried under sterile conditions. Cells (0.8-1 x 10^6^) were seeded on the dried coverslips and allowed to attach for at least 1 hr. After adhering to the coverslips, the cells were washed briefly with PBS at RT, fixed in ice cold 2% formaldehyde (FA, Polysciences, Cat. No. 04018-1) in PBS for 7.5 min and permeabilized in 1% FA and 0.25% Triton X-100 (Sigma, Cat. No. T8787) in PBS for 7.5 min on ice. Following two washes with ice-cold PBS, cells were blocked in 3% BSA (Serva, Cat. No. 11930.03) in PBS for 1 hr at RT. Primary antibodies were diluted in 0.1% Triton X-100 and 1.2% normal donkey serum (Johnson Immuno, Cat. No. 017-000-121) in PBS and incubated overnight at 4°C. After two PBS washes at RT, cells were incubated with secondary antibodies diluted in 0.1% Triton X-100 and 1.2% NDS in PBS for 1 hr at RT. For more information on the antibodies and fluorophores used, see Supplementary Table 1. Following two PBS washes, nuclei were stained with DAPI (Sigma, D-9542) in PBS for 2 min. Coverslips were washed once with PBS and once with water, then mounted using Vectashield antifade mounting medium for fluorescence (Vector Labs, Cat. No. H-1000) and sealed with nail polish.

*Confocal microscopy* was performed at the Core Facility Bioimaging of the Biomedical Center with an inverted Leica SP8X WLL microscope, equipped with 405 nm laser, WLL2 laser (470 - 670 nm) and acusto-optical beam splitter. Images were recorded with a 100x/1.4 NA STED white objective with zoom factor of 4.5, yielding a field of view of 25.8 × 25.8 μm. The image format was set to 512 × 512 pixels, resulting in a pixel size of 50.4 nm. Z-stacks were acquired in xyz scan mode using galvo-based z-scanning (z-galvo) with a constant step size of 0.24 μm and unidirectional x-scanning at a speed of 200 Hz (pixel dwell time: 7.7 µs). The confocal pinhole was set to 1.0 Airy units using 580 nm as reference wavelength. Fluorescence signals of DAPI, MSL2 (Cy2), MSL3 (Cy3) and H4K16ac (Alexa Fluor 647) were recorded frame sequentially to avoid channel crosstalk with the following excitation and detection settings (excitation wavelength; detector: detection window): DAPI: (405 nm; PMT1: 415 – 480 nm), Cy2: (488 nm; HyD2: 498 – 535 nm), Cy3: (551; HyD4: 561 - 620 nm), Alexa Fluor 647 (650 nm; HyD5: 660 – 710 nm). Detector gain was set to 700 Volt (PMT1) and 50% (HyDs in standard mode). Images were acquired at 8-bit depth and subsequently deconvolved using the Huygens Software (v17.04) and stored at 16-bit depth (512 × 512 × 20 z-planes; z-step ∼0.24 µm). Maximum intensity projections and scaling of image intensity was performed using Fiji (ImageJ, 2.16.0) (39).

*Widefield microscopy* was performed at the Core Facility Bioimaging of the Biomedical Center with an inverted Leica THUNDER Imager 3D Live Cell TIRF with DMi8 stand equipped with LED5 multi-LED illumination (390/475/560/635 nm) and fast filter wheel with emission filters centered at 460, 535, 590, and 642 nm. Images were recorded with an HC PL APO 100×/1.47 oil CORR TIRF objective, yielding a field of view of 133 × 133 μm per tile (2048 × 2048 pixels at ∼65 nm/pixel sampling, Leica K8 sCMOS camera, 16-bit depth). Z-stacks were acquired sequentially across 4 channels using ∼0.3 μm z-steps and a region spanning 3×4 fields of view. Images were acquired with the following settings: DAPI for DNA (Ex 391/32 nm, Em 435/30, 8% LED, 100 ms), Cy2 for MSL2 (Ex 479/33 nm, Em 519/25 nm, 12% LED, 100 ms), Cy3 for MSL3 (Ex 554/24 nm, Em 594/32 nm, 15% LED, 300 ms), and Alexa Fluor 647 for H4K16ac (Ex 638/31 nm, Em 695/58 nm, 20% LED, 200 ms). Gain was set to 1 (HDR Combined Gain-16 bit mode) and aperture diaphragm to 8 for all channels. Further processing of the images was performed using Fiji and macros scripts.

#### Image processing

Images were processed (maximum intensity projections and scaling) uniformly across conditions for display purposes. Image analysis was performed in R (v4.5.1) with the following packages: EBImage (v4.50.0, for image processing and feature extraction)(40), tidyverse suite of tools (v2.0.0, for preprocessing, statistical analysis and visualization)(41), corrr (v0.4.5, correlation analysis), cluster (v2.1.8.1, clustering validation) and umap (v0.2.10.0, dimensionality reduction)(42). Briefly, single-cell images were read as multi-channel TIFF files, and the MSL2 channel (channel 2) was extracted for analysis. Images were normalized to a 0–1 intensity range prior to feature extraction. For image-based profiling, MSL2 immunofluorescence intensity distributions and Haralick texture features were extracted from single cells (43). For each cell, 46 quantitative features were extracted. For segmentation, images were Gaussian-blurred and thresholded at 0.85 to identify high-intensity regions corresponding to MSL2-enriched foci. Prior to dimensionality reduction, highly correlated variables were removed. Uniform Manifold Approximation and Projection (UMAP) was performed using the umap R package with parameters n_neighbors = 100 and min_dist = 0.1. Statistical differences in feature values between clusters were assessed using Kruskal-Wallis tests with effect size filtering (ε² > 0.1, FDR-adjusted p < 0.05), followed by pairwise Wilcoxon rank-sum tests with FDR correction to identify cluster-specific features. The proportion of cells assigned to each cluster was quantified for each genotype and rescue condition. Data for one out of four independent replicates are shown, a cumulative UMAP colored by genotype for each other replicate can be found in Supplementary Fig. 1A.

#### Chromosome spreads and karyotype analysis

Chromosome spreads were prepared from Schneider S2 cells as described in (44), except that cells were fixed for 10 min using 3.7% formalin (Merck, 8187081000) in PBST (PBS+0.1% Triton X-100) solution and only stained with 1 μg/ml DAPI (4’, 6-diaminidino-2-phenylindole) in PBST. Slides were imaged using 100x oil immersion objective on Nikon-CREST widefield microscope. Image deconvolution was performed with the Huygens Essential Software from SVI (version 24.10.0p7 64b). Individual chromosome identification was done using the *Drosophila* S2 cell karyotype in Figure S3a. of Olszak et al. (2011) (44). For a comprehensive summary of the mitotic spreads analyzed in this study see Supplementary Table 2.

### Data analysis

#### Sequencing data preprocessing

Paired-end CUT&RUN and ChIP-MNase sequencing reads were aligned to the *Drosophila melanogaster* dm6 reference genome (UCSC) using Bowtie2 (v2.2.9, --local --very-sensitive-local --no-unal --no-mixed --no-discordant)(45). Maximum fragment sizes were set to 700 bp for CUT&RUN and 1000 bp for ChIP-MNase to accommodate the distinct fragment size distributions of each method. SAM files were converted to BAM format retaining high-quality alignments (samtools v1.22.1, samtools view -q 2)(46), name-sorted, and converted to BED format using BEDtools (v2.31.1)(47) bamToBed to extract fragment coordinates. CUT&RUN data were normalized to total read counts. Scaling factors were calculated as 10^6^ divided by the total number of filtered reads per sample (excluding secondary, supplementary, and unmapped reads, and reads with mapping quality < 30). Normalized coverage tracks were generated using deepTools2 (v3.5.0)(48) bamCoverage at 10 bp resolution with target-specific smoothing (20 bp for MSL2/PPI; 80 bp for H4K16ac/IgG). Pairwise target-to-control comparisons were computed using bamCompare to generate ratio tracks with a pseudocount of 1. Pre-immune serum (PPI) and rabbit IgG were used as negative controls for the targets MSL2 and H4K16ac respectively. For ChIP-MNase data, spike-in normalization was performed using reads aligned to the *Drosophila virilis* droVir3 genome. Scaling factors were calculated as 10^6^ divided by the spike-in read count for each sample. Normalized to Input coverage tracks were generated using BEDtools (v2.31.1) genomecov and converted to bigWig format using bedGraphToBigWig (UCSC utilities, 3.4.1)(49). Biological replicates were merged (n=3 or n=2 for stable lines) using Wiggletools (1.2.2)(50) and visualized using the IGV genome browser (v2.19.4)(51) and the dm6 *Drosophila melanogaster* genome release. For correlation analysis between the CUT&RUN biological replicates across targets and genotypes, after total reads and negative control normalization see Supplementary Fig. 2A.

#### Window-based binding site identification using csaw

DCC binding sites were identified from CUT&RUN data derived from wild-type cells using a window-based differential binding approach implemented in the csaw R/Bioconductor package (v1.42.0,(52)). Analysis parameters were optimized separately for the transcription factor MSL2 and the histone modification H4K16ac. For MSL2, reads were counted in 20 bp sliding windows with fragment extension of 100 bp and maximum fragment length of 250 bp. For H4K16ac, reads were counted in 120 bp sliding windows with fragment extension of 150 bp and maximum fragment length of 500 bp; windows were retained if signal was present in at least three replicates. Read counts were TMM-normalized (edgeR, v4.6.3)(53), and dispersions were estimated robustly. Differential binding between target and negative control was assessed using quasi-likelihood F-tests. Significance thresholds were FDR ≤ 0.01 and |log2FC| ≥ 2 for MSL2, and FDR ≤ 0.05 and |log2FC| ≥ 1 for H4K16ac. Thresholds were chosen to reflect the biology of each target: stringent parameters for MSL2 to identify discrete, high-confidence binding sites, and more permissive parameters for H4K16ac to capture the broader landscape of histone acetylation across MSL-bound domains, while maintaining optimal signal-to-noise ratios for each dataset. Significant windows were merged within 50 bp to define MSL2 CUT&RUN csaw sites (MCCS) and H4K16ac CUT&RUN csaw sites (HCCS). Significant windows passing all filters were merged into contiguous binding regions using GenomicRanges package (v1.60.0)(54) with a maximum gap of 50 bp. Merged regions were annotated to genomic features using the ChIPseeker package (v1.44.0)(55) with TxDb.Dmelanogaster.UCSC.dm6.ensGene (v3.12.0), defining promoters as 1 kb upstream from the TSS.

#### Analysis of CUT&RUN data

High resolution merged MSL2 binding sites (MCCS: MSL2 CUT&RUN Csaw Sites) were compared to previously characterized High Affinity Sites (HAS) (29) and Pioneering on the X (PionX) sites (56). Overlap analysis was performed by creating a union of all peak sets using GenomicRanges, then determining membership of each unified region in the original sets using countOverlaps. Overlaps were visualized using UpSetR (v1.4.0)(57). Signal intensity at binding sites was quantified by extracting mean enrichment values from merged bigWig files (ratio of target/control) using rtracklayer (v1.68.0)(58).

#### Heatmap and Metaplot Visualization

Signal profiles at binding sites were visualized using deepTools2 (v3.5.0). For site-centered analysis, computeMatrix reference-point was used with flanking 2 kb windows centered on binding site midpoints. For gene body analysis, computeMatrix scale-regions was applied to MSL2-bound genes with 2 kb upstream/downstream flanks and 5 kb scaled gene body regions. Matrices were sorted by mean signal in descending order using the WT sample. Heatmaps were generated with plotHeatmap using color maps with z-score ranges of 0-4 and 0-8.

#### Motif Analysis

Genomic sequences underlying DCC binding sites were extracted from the Drosophila melanogaster genome (dm6) using the BSgenome.Dmelanogaster.UCSC.dm6 R package (v1.4.1). Binding site regions (MCCS, HAS, PionX) were resized to 200 bp centered on the summit, and sequences were extracted using the Biostrings package (v2.76.0)(59). Control background sequences were generated by randomly sampling genomic regions matched for chromosome composition and sequence length. For each binding site, a corresponding background region was selected from the same chromosome with matching width. Background regions overlapping true binding sites were excluded to ensure specificity. Random sampling was performed with a fixed seed (42) for reproducibility. STREME (Sensitive, Thorough, Rapid, Enriched Motif Elicitation; MEME Suite v5.5.8) (60)was used for discriminative motif discovery comparing target sequences against the matched background set. Parameters: --dna --nmotifs 5 --minw 6 --maxw 20 --thresh 0.05. STREME was applied to all binding site categories. Discovered motifs were compared to known *Drosophila* transcription factor binding motifs using Tomtom (MEME Suite v5.5.8)(61) against multiple databases: FlyReg v2, Fly Factor Survey, OnTheFly 2014, and dmmpmm2009. Parameters: -min-overlap 5 -dist pearson -evalue -thresh 10.0.

#### DNA copy number estimation from ChIP-MNase input samples

Chromosome copy number was estimated from ChIP-MNase-seq input samples, which represent chromatin that has been mildly sonicated, MNase-digested, and pre-cleared (see ChIP-MNase sequencing above), and thus serves as a proxy for genomic DNA content. Replicate input BAM files were processed to generate read coverage across 50 kb non-overlapping genomic bins. For each bin, read counts were converted to an RPKM-like metric (reads per kilobase per million mapped reads) to normalize for bin length and library size differences. Spike-in normalization was applied to account for technical variation between samples: each sample was aligned to both the *Drosophila melanogaster* dm6 reference genome and the *D. virilis* spike-in genome (droVir3). The ratio of D. melanogaster to spike-in reads was used to derive per-sample scaling factors, enabling quantitative comparison of absolute DNA content across conditions. Downstream processing, statistical analyses, and visualizations were performed in R (v4.5.1) using the GenomicRanges (v1.60.0), Rsamtools (v2.24.1)(62), and tidyverse (v2.0.0) packages.

#### Chromosome-level copy number analysis

Per-chromosome copy numbers were calculated by computing the median of bin-level copy number values for each chromosome arm (chr2L, chr2R, chr3L, chr3R, chr4, chrX) within each biological replicate. To convert relative coverage values to absolute copy numbers, values were scaled to the wild-type autosomal baseline: the median of per-chromosome medians across autosomes (chr2L, chr2R, chr3L, chr3R) in wild-type samples was set to the expected copy number of 4, reflecting the tetraploid nature of S2 cells. All chromosome values were then scaled proportionally using this baseline. For each condition (WT, KO-A, KO-B, Rescue), the mean and standard deviation of scaled copy numbers were calculated across biological replicates (n = 3 per condition). Error bars in figures represent the standard deviation of replicate-level medians converted to the absolute copy number scale. A comprehensive comparison of the scaled copy number per bin for the X chromosome, for all conditions (including replicates) can be found in Supplementary Fig. 4.

### RNA sequencing and analysis

#### RNA isolation and library preparation

Total RNA was isolated from S2 cells using the RNeasy Mini Kit (Qiagen, Cat. No. 74106) according to the manufacturer’s instructions. Cells were counted, and approximately 3–6×10^6^ cells were harvested per sample. RNA concentration was measured, and samples were stored at −80°C. RNA was treated with DNase as described earlier. DNase-treated RNA was cleaned-up using the RNeasy Mini Kit and quality-assessed by Qubit RNA quantification and TapeStation analysis to verify RNA integrity. Strand-specific RNA-seq libraries were prepared from 100 ng of DNase-treated total RNA using the QIAseq FastSelect RNA Library Kit (Qiagen, Cat. No. 334302) according to the manufacturer’s protocol with the following modifications: RNA fragmentation was performed at 95°C for 10 minutes and rRNA depletion was performed using 0.1× FastSelect reagent. Libraries were amplified for a total of 20 PCR cycles, and concentrations were measured by Qubit HS DNA assay. Library size distribution was assessed by TapeStation, and a size selection step was performed using QIAseq beads (Qiagen, Cat. No. QC13.3) (0.6× ratio for removal of larger fragments, followed by 1× for cleanup) to remove fragments with aberrant size distributions. Final libraries were eluted in 10 μl nuclease-free water and submitted for sequencing. RNA-seq was performed in two batches. Batch 1 included wild-type S2 cells (n=4 biological replicates) and *roX2* knockout clone KO-B (n=4 biological replicates). Batch 2 included additional wild-type replicates (n=2), *roX2* knockout clone KO-A (n=4 biological replicates), and rescue cells expressing full-length *roX2* (res; n=2 biological replicates). Biological replicates were derived from independently passaged cell populations.

#### RNA-seq data processing and analysis

Raw paired-end sequencing reads were processed using the snakePipes (v3.2.0) noncoding RNA-seq workflow (63). Adapter sequences were trimmed using cutadapt via Trim Galore (v4.6/0.6.10). Quality control was performed using FastQC (v0.12.1)(64) and summarized with MultiQC (1.25)(65). Trimmed reads were aligned to the *Drosophila melanogaster* dm6 reference genome using STAR (v2.7.10b)(66) with parameters optimized for multi-mapping reads (--outFilterMultimapNmax 1000 --outFilterMismatchNoverLmax 0.1 --outSAMstrandField intronMotif). Gene-level read counts were quantified using the Ensembl gene annotation (release 96). Gene expression was quantified as transcripts per million (TPM) for visualization and as raw counts for normalization. RNA-seq differential expression analysis was performed using DESeq2 (v1.48.2) (67) on all annotated genes. Raw read counts were filtered to retain genes with at least 10 counts in a minimum number of samples equal to the smallest experimental group size. Differential expression was assessed using a batch-aware design with the Wald test. Genes were considered differentially expressed (DE) if they met the following criteria: adjusted p-value (padj) < 0.01 and |log2 fold-change| > 1. Pairwise comparisons were performed between each knockout condition, rescue and wild-type (WT). Volcano plots were generated to visualize DE genes, with the top 20 genes by significance and top 20 by absolute log2 fold-change annotated, Supplementary Fig. 3C.

#### Gene expression and dosage compensation analysis

For visualization of individual gene expression (e.g., *roX1* and *roX2*), TPM values were log10-transformed (log10(TPM + 1)) and plotted per condition with individual replicates shown. Error bars represent the standard deviation of log10-transformed values across replicates. To assess dosage compensation status, analysis was restricted to protein-coding genes located on major chromosome arms (chr2L, chr2R, chr3L, chr3R, chr4, chrX). Genes with fewer than 10 counts in at least 3 samples (or 2 when not available) were excluded. Raw counts were normalized using DESeq2 median-of-ratios normalization. For each knockout condition, mean normalized counts were calculated across replicates, and log2 fold changes relative to wild-type were computed per gene as log2((KO + 1) / (WT + 1)). Log2 fold change distributions were compared between X-chromosomal and autosomal genes for each genotype to evaluate whether X-linked gene expression was selectively affected by *roX2* loss.

#### Copy number-corrected expression analysis

To assess the relative contributions of DNA copy number and transcriptional regulation to X chromosome expression, copy number-corrected expression values were calculated using R with DESeq2 (v1.48.2), tidyverse (v2.0.0), and patchwork (v1.3.2)(68) packages. For each condition, per-gene RNA-seq expression values (DESeq2-normalized counts) were divided by the corresponding chromosome copy number estimated from ChIP-MNase sequencing input samples. X:A (X/Autosome) expression ratios were calculated both with and without copy number correction to distinguish canonical DCC-mediated dosage compensation from other mechanisms. This method enables the decomposition of dosage compensation into copy number and transcriptional contributions. The buffering effect of elevated X chromosome copy number was quantified as the ratio of observed X:A expression to the expected ratio based on copy number alone.

## Results

### Generation and validation of *roX2* knockout S2 cell lines

We sought to investigate the functional requirements of the long non-coding *roX* RNA for dosage compensation in *Drosophila* in a simple, proliferative cell model. Descendants of the S2 cell line have proven very useful for studying dosage compensation. These aneuploid cells are mainly tetraploid, having two X chromosomes, but four copies of each autosome. They compensate the 2-fold reduced number of X chromosomes relative to autosomes transcriptionally through about equal contributions of a generic feedback mechanism that works on all chromosomes and a fixed 1.4-fold transcription boost due to the action of the MSL dosage compensation complex (DCC)(28). The role of the *roX* RNAs had not been analyzed in this well-controlled system.

S2 cells do not express the *roX1* RNA to significant levels and dosage compensation in these cells thus is effectively mediated by *roX2* (14). To generate *roX*-deficient cell lines, we deleted the *roX2* gene using CRISPR–Cas9 genome editing (Fig. 1). We transfected S2 cells with plasmids expressing Cas9 and encoding three sgRNAs targeting the *roX2* locus. Single clones harboring the plasmids were selected with blasticidin, isolated and expanded in a time course of 6 weeks (Fig. 1A). Only two independent knockout clones were recovered, designated KO-A and KO-B, and validated by PCR and sequencing of deletion junctions. Genome-browser visualization of DNA sequencing reads from ChIP input samples (see below) across the *roX2* locus verified the deletion of the *roX2* region in both alleles (Fig. 1B).

**Figure 1.**
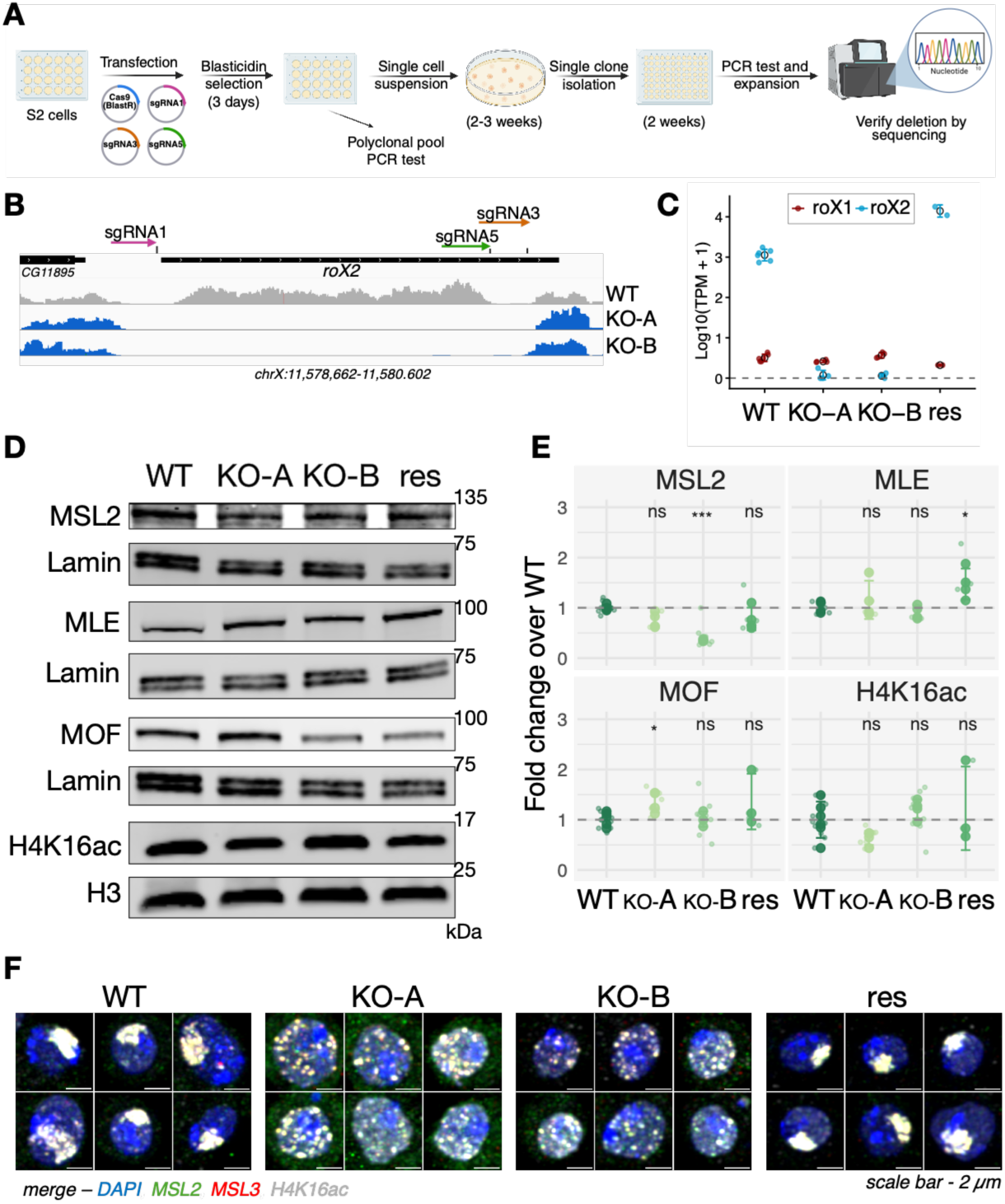
Generation and characterization of *roX2* knockout S2 cell lines. (A) CRISPR–Cas9 strategy and workflow used to isolate *roX2* knockout clones. (B) Deletion of the *roX2* gene. Top, schematic illustration of the positions of three sgRNAs relative to the *roX2* gene (black line). Bottom, genome-browser view of ChIP–MNase input coverage tracks across the *roX2* locus (coordinates below) for wild-type (WT, grey) and the two *roX2* knockout clones (KO-A and KO-B, blue). Loss of coverage reveals the deletion of the *roX2* region in both clones. (C) Quantification of *roX* RNAs in wild-type, KO and rescue (res) lines. Total RNA-sequencing (RNA-seq) reads [log10(TPM)+1 values] for *roX1* (blue) and *roX2* (red) are plotted for the indicated lines. Data points denote biological replicates. *roX2* expression is undetectable in KO-A and KO-B and restored in the rescue line; *roX1* expression remains very low across conditions. (D) Representative immunoblot probing DCC proteins and H4K16 acetylation (H4K16ac). Whole-cell extracts from WT, KO-A, KO-B and rescue cells were probed for MSL2, MLE, MOF, and H4K16ac. Lamin and histone H3 serve as loading controls. Positions of size markers (kDa) are indicated. (E) Quantification of immunoblot signals from Westerns as in (D). Signals are shown as fold change relative to WT after normalization to Lamin (non-histone proteins) or histone H3 (H4K16ac). Each data point represents an independent biological replicate; technical replicates are shown as smaller points, when applicable. The dashed line indicates WT protein expression (fold change=1). Statistical significance is annotated as indicated (ns, not significant). (F) Confocal microscopy to visualize DCC enrichment on the X chromosome. Galleries of six representative confocal images of nuclei of the indicated cell lines, stained for MSL2 (green), MSL3 (red) H4K16ac (grey) and DAPI (blue) marking nuclei. The images are deconvolved Z-maximum projections of multiples stacks; channels are merged and auto-scaled for display purposes. WT cells show the characteristic enrichment of MSL2/MSL3 and H4K16ac at the X-chromosome territory. In knockout cells the complex is delocalized. The rescue line restores X association. Scale bar, 2 μm.

RNA sequencing and qPCR analysis confirmed the absence of *roX2* transcripts in both KO-A and KO-B (Fig. 1C, Fig. 2B, and Supplementary Fig. 3C). *RoX1* expression remained negligible as in wild-type (WT) cells, indicating that *roX1* is not increased to compensate for the loss of *roX2*. In a functional rescue experiment, we generated a stable line expressing a full-length *roX2* transgene in the KO-B background (res), which restored *roX2* transcripts to levels comparable to WT (Fig. 1C and Supplementary Fig. 3C).

**Figure 2.**
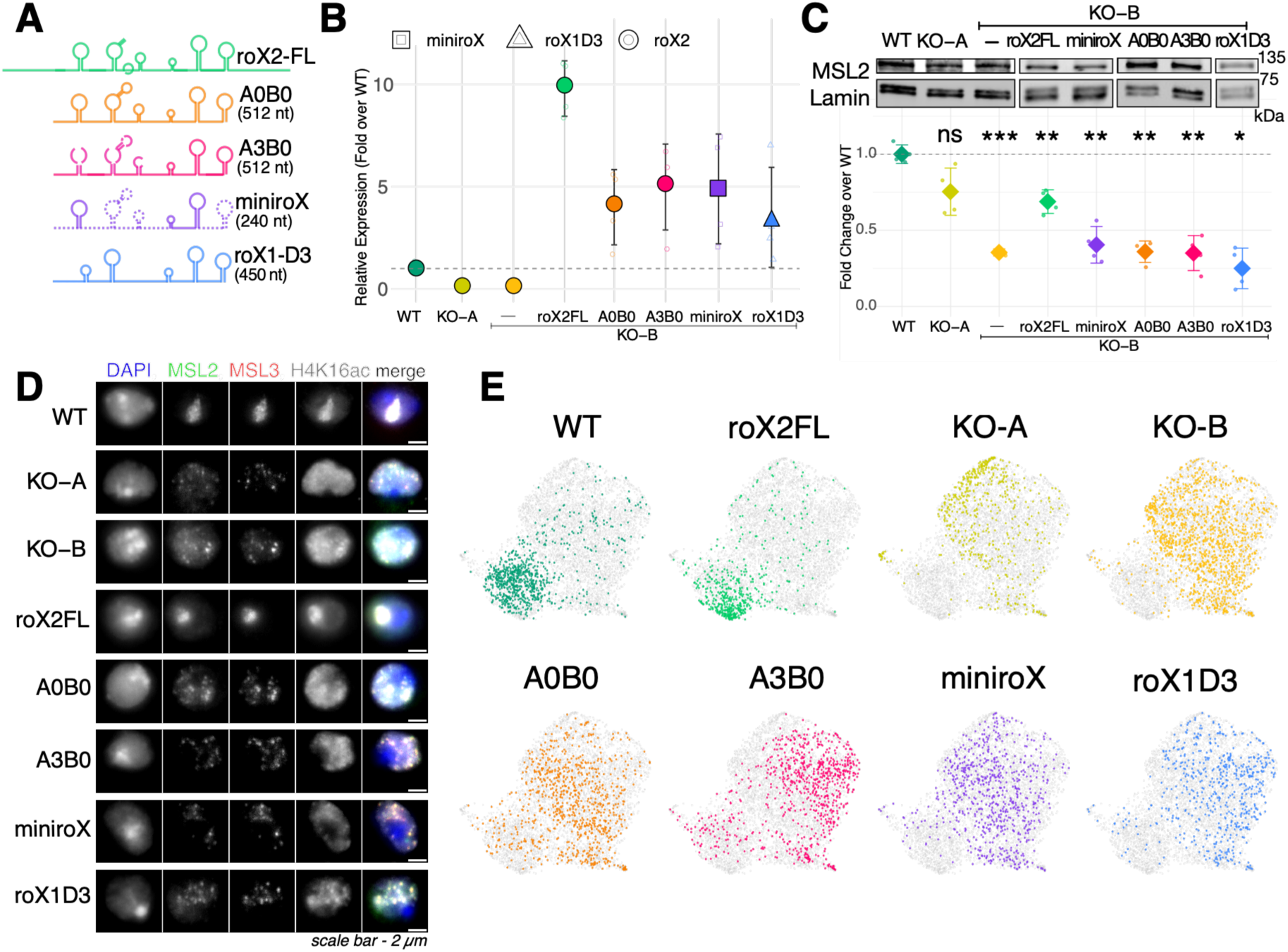
An experimental system to study *roX* requirements for X chromosome targeting. (A) Schematic depiction of *roX* constructs used for rescue experiments. Conserved stem–loop elements are indicated. RNA lengths are shown in parentheses. For details see text. (B) Expression of *roX* transgenes. RT–qPCR quantification of *roX* variant expression in stable cell populations using specific primer sets for *roX2* (circles), *miniroX* (squares), and *roX1-D3* (triangles). Values are normalized to *GAPDH* RNA and expressed relative to WT using the ΔΔCp method. The mean ± SD of 4 biological replicates is shown. The dashed line indicates WT expression. (C) MSL2 protein levels in cells expressing *roX* variants. Top, representative immunoblot of MSL2 across WT, KO clones and rescue lines expressing the indicated constructs. Lamin serves as loading control. Bottom, quantification of MSL2 normalized to the internal control and relative to WT expression (dashed line). The mean signal of 4 biological replicates (data points) ± SD is shown. Statistical significance is indicated as annotated (ns, not significant). (D) Widefield microscopy images of representative cells illustrating DCC localization and H4K16ac upon *roX* RNA expression. Cells were stained for MSL2 (green), MSL3 (red) H4K16ac (grey) and DAPI (blue) marking nuclei. Individual channels are displayed in greyscale. All images were scaled uniformly for display. Scale bar, 2 μm. (E) Classification of MSL2 localization states in single cells by image-based profiling. UMAP projection of single cells derived from widefield microscopy and quantified using MSL2 immunofluorescence intensity distributions and Haralick texture features. Each point represents one cell, colored by genotype/condition, as indicated. One representative replicate of this experiment is shown, for UMAP projections of the other replicates see Supplementary Fig. 1A).

The MSL proteins and *roX* RNA are stable as components of the DCC complex, but unstable if the integrity of the DCC is disrupted by depletion of a protein subunit (21,69). To explore whether *roX2* loss affected the MSL protein abundance, we monitored MSL protein levels in the KO lines by immunoblotting. We found that MSL2 was moderately reduced in both KO clones, but more prominently in KO-B, while MLE and MOF levels remained largely unchanged (Fig. 1D, E). Re-expression of *roX2* in the rescue line partly restored wild-type MSL2 levels. The levels of H4K16ac, the epigenetic readout of DCC action, showed modest variation but overall were not substantially altered.

The hallmark of functional dosage compensation in male *Drosophila* cells is the selective binding of DCC to the X chromosome. Immunostaining of the MSL proteins visualizes their binding to a compact chromosomal territory. Depletion of any MSL protein abolishes the X chromosome ‘territory’ staining of the complex (21,27,70). Confocal microscopy of WT cells demonstrated the characteristic co-localization of MSL2, MSL3 and H4K16ac at the dosage-compensated X chromosome (Fig. 1F). Strikingly, this subnuclear enrichment was lost in both *roX2* knockout clones, where the complex was delocalized throughout the nucleus. Expression of the *roX2* transgene in the rescue line restored X-chromosome association of the DCC, demonstrating that *roX2* is required for faithful targeting.

### An experimental system for *roX* structure-function analyses

The successful rescue of DCC function by expression of *roX2* in the KO-B cell population suggested a strategy through which features of *roX* RNA function could be evaluated. As a proof of principle, we tested several *roX2* derivatives that in the systematic analyses of Akhtar and colleagues had been shown to support a certain level of viability during male fly development. Their *A0B0* construct ((17), Fig. 2A) represents ‘exon 3’ of *roX2* with a total length of 512 bp, which contains all known binding sites for the helicase MLE. A derivative of this sequence, designated *A3B0,* has the three 5’ stem-loop (SL) structures mutated that attract MLE in an ATP-independent manner, but maintains critical 3’ SLs that are the target for MLE remodeling (17,19,20). Ilik et al. also generated a ‘*minirox’* variant, in which central *roX2* sequences were deleted, leaving the 5’ and 3’ SLs intact (18). Their data suggest that this might be the shortest *roX* derivative that can sustain male life. We selected the *A0B0*, *A3B0* and *miniroX* constructs to explore their ability to complement the *roX*-deficiency in KO-B cells. Finally, we tested the hypothesis that the long ‘*roX1’* RNA contains a *roX2*-like module at its 3’ end. The 3’ *roX1-D3* region resembles *roX2* structurally [SL structures occluding ‘*roX* boxes’ (17)]. Since *roX1* is functionally redundant with *roX2* for male viability, it is possible that the 3’ portion of *roX1* represented in *roX1-D*3 can substitute for *roX2* in our system.

We generated stable cell populations expressing *roX2* sequences corresponding to *A0B0*, *A3B0, miniroX*, and *roX1-D3*. RT–qPCR analysis confirmed expression of all transgenes at 3-5-fold elevated levels compared to endogenous *roX2* in WT cells (Fig. 2B). Immunoblot analyses revealed that expression of full-length (FL) *roX2* partially restored MSL2 levels, whereas none of the test constructs improved MSL2 levels (Fig. 2C). We assessed DCC localization by widefield microscopy, staining for MSL2, MSL3 and H4K16ac (Fig. 2D). FL *roX2* restored the characteristic X-chromosome territory enrichment pattern observed in WT cells. In contrast, expression of the truncated roX variants resulted in diffuse staining patterns lacking clear territory enrichment.

To assess the rescue capacity in an unbiased manner, we employed image-based profiling of single cells using MSL2 immunofluorescence intensity distributions and textural properties, followed by UMAP dimensionality reduction (Fig. 2E and Supplementary Fig. 1A). Unsupervised clustering identified two broadly distinct populations of cells. The WT cells (dark green) occupied a discrete region at the lower left of the UMAP space, whereas KO-A and KO-B cells were assigned to clearly separable areas within a larger cluster. Consistent with their restored territory staining, cells expressing FL *roX2* predominantly co-localized with the WT cluster. In contrast, cells expressing *A0B0*, *A3B0*, *minirox*, or *roX1-D3* failed to co-localize with the WT population. Minor fractions of cells expressing *roX* derivatives *A0B0* and *A3B0* were mapped to the ‘territory’ area of the UMAP, predominantly occupied by WT and FL *roX2*.

*Minirox* and *roX1-D3* remained clearly distinct. Interestingly, *A0B0* and *A3B0* occupied partially overlapping but distinguishable patterns in the UMAP, indicating that the texture-based analysis could potentially distinguish subtle differences in DCC localization patterns. Independent repetitions of this experiment with distinct biological replicates showed similar results (Supplementary Fig. 1A). CUT&RUN in the stable transgenic cell lines confirmed the widefield microscopy results, namely constructs that partially rescue DCC targeting (*roX2*-FL and *A0B0*) show some degree of X-chromosome enrichment, whereas non-rescuing constructs (*A3B0, miniroX, roX1-D3*) do not support X-chromosome targeting (Supplementary Fig. 1B).

### Genome-wide mapping of DCC occupancy by CUT&RUN

The confocal images showed that the DCC was delocalized from the X chromosome in the absence of *roX*. To characterize DCC binding at a higher resolution genome-wide, we performed CUT&RUN for MSL2 and H4K16ac in WT and *roX2* KO cell lines (Supplementary Fig. 2A). We had shown earlier that CUT&RUN allows to identify a larger number of X-chromosomal binding sites compared to ChIP (25). For a direct comparison we mapped MSL2 by CUT&RUN and ChIP in WT, KO and rescue lines. Representative genome-browser views of sites of MSL2 and H4K16ac localization on the X chromosome are shown in Figure 3A. The comparison with matched ChIP data reveals that CUT&RUN indeed identifies more MSL2 binding sites. The ChIP peaks are much broader, presumably because of variable crosslinking of MSL2 to neighboring chromatin (12). For the H4K16 acetylation, the correspondence between the two methods is excellent. This is also evident when we compare the signal of MSL2 and H4K16ac over gene bodies at MSL2-bound genes (Supplementary Fig. 2B, C).

**Figure 3.**
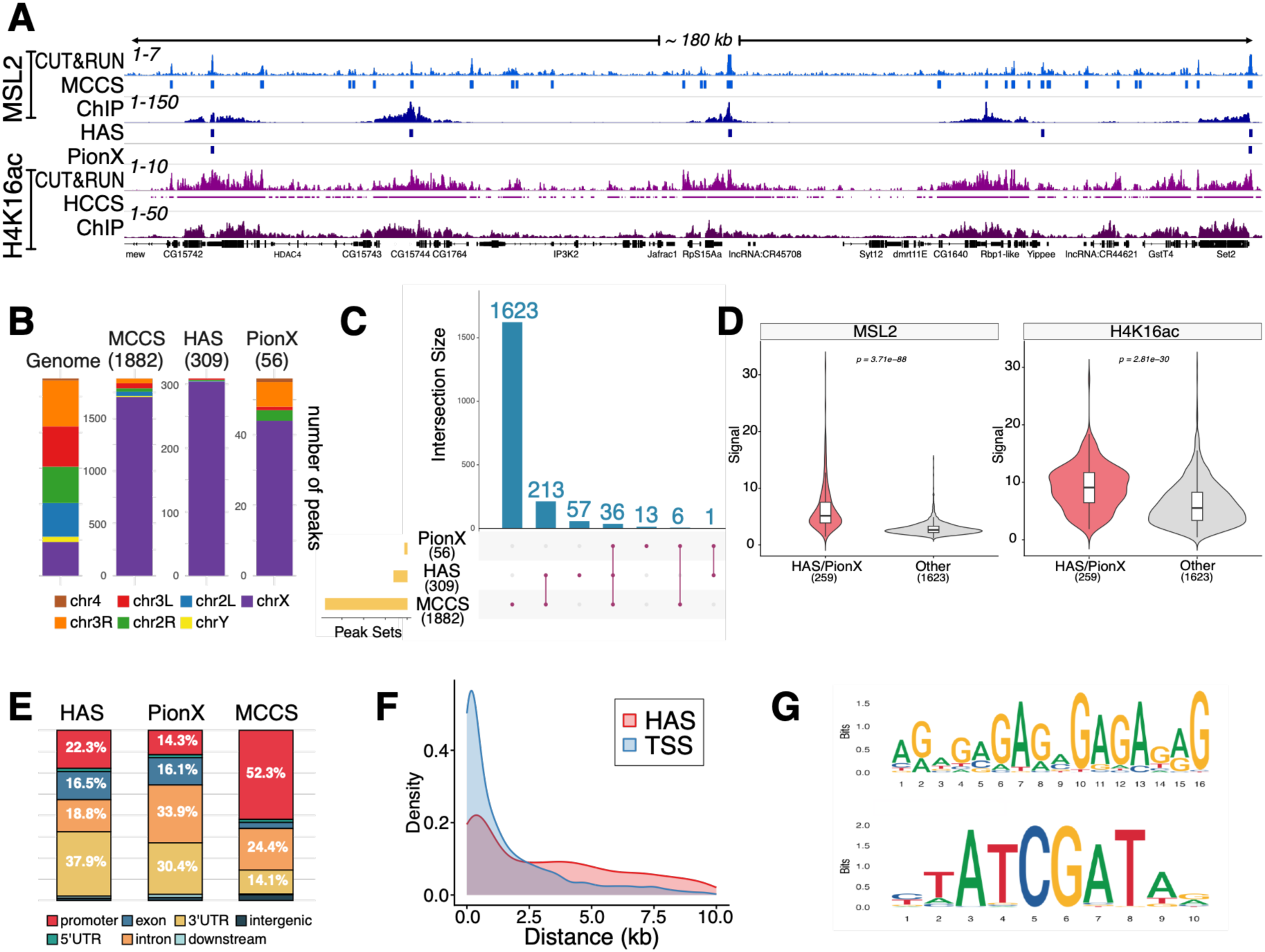
Genome-wide mapping of DCC occupancy by CUT&RUN and csaw. (A) Representative genome-browser views of merged MSL2 and H4K16ac CUT&RUN profiles (n = 3, Supplementary Fig. 2A) across a ∼180 kb region of the X chromosome, along with matched ChIP-MNase sequencing tracks (n = 3). Canonical DCC binding site annotations (HAS and PionX) and newly identified CUT&RUN peak sets computed by csaw are indicated (MCCS: MSL2 CUT&RUN csaw sites; HCCS: H4K16ac CUT&RUN csaw sites). Track scales are shown; gene names are displayed below. (B) Chromosomal distribution of canonical (HAS, PionX) and novel (MCCS) DCC binding sites. Bars indicate the number of sites assigned to each chromosomal arm and the distribution of peaks among the chromosomes in a color code. The number of peaks in each category is provided in parentheses. (C)UpSet plot showing the overlap between MCCS and previously mapped HAS and PionX signature MSL2 binding sites. (D) CUT&RUN signal intensities at MCCS that overlap (HAS/PionX, red) or not (Other, grey) with HAS/PionX signature sites. Aggregate MSL2 and H4K16ac coverages at classified MCCS sites are shown. Numbers of sites in each group are indicated in parentheses. P-values: Wilcoxon rank-sum tests. (E) Genomic feature annotation of MSL2 binding sites. The bar graph shows the fraction of HAS, PionX and MCCS sites assigned to genomic features indicated by the color code. Approximately half of MCCS map to promoter regions (red). (F) Proximity of MCCS to transcription start sites (TSS) and HAS. Density distributions showing distances from MCCS to the nearest TSS and to the nearest HAS, as indicated. (G) Sequence logos of motifs enriched at MCCS-associated regions. The top scoring logo corresponds to the extended MSL recognition element (MRE). The bottom motif resembles DREF (TATCGATA) or BEAF-32 binding sites (ATCGAT). Letter height denotes information content.

To capture the unique characteristics of CUT&RUN more accurately than traditional peak calling algorithms, we applied a window-based differential binding analysis (csaw)(71). Our approach counts reads in sliding windows across the genome and identifies regions with statistically significant enrichment over background. Using stringent thresholds and optimized parameters for each target (see Methods), we identified statistically enriched regions and merged proximal windows to define MSL2 CUT&RUN csaw sites (MCCS) and H4K16ac CUT&RUN csaw sites (HCCS).

This analysis identified 1,882 MCCS, with approximately 90% mapping to the X chromosome (Fig. 3B). As expected, MCCS encompassed many previously characterized High Affinity Sites (HAS, 309 sites) and PionX (56 sites) signature sites (Fig. 3A). However, UpSet plot analysis revealed that 1,623 MSL2 binding sites, 86% of all MCCS, had not been described previously (Fig. 3C), indicating that ChIP predominantly detects only the highest-occupied sites. Consistent with this, the HAS and PionX signature sites within MCCS displayed significantly higher MSL2 and H4K16ac signals compared to novel MCCS (Wilcoxon rank-sum test, p < 0.001; Fig. 3D), confirming a hierarchical binding landscape where previously known sites represent the most robustly occupied DCC targets (Fig. 3D).

Genomic feature annotation revealed that approximately half of MCCS mapped to promoter regions (within 1 kb upstream of transcription start sites, TSS). In contrast, most of HAS and PionX sites (about 73% and 80%, respectively), are localized to gene bodies (Fig. 3E). Most of the newly identified MSL2 binding sites map to promoter regions, as visualized by plotting the distances of MCCS to TSS as compared to HAS (Fig. 3F). Motif discovery analysis at MCCS peaks found the extended MSL recognition element (MRE) as the top-scoring motif that had been identified in previous studies (25,56,72–74) (Fig. 3G). Interestingly, a palindromic motif that to our knowledge has not yet been implicated with dosage compensation scored as second-best (consensus: YTATCGATAR; E-value = 4.8×10⁻⁴). Search in the Tomtom database revealed highly significant matches to BEAF-32 (Boundary Element-Associated Factor; p-value = 3.47×10⁻⁷, E-value = 4.93×10⁻⁴) and DREF (DNA replication-related element factor; p-value = 2.25×10⁻⁵). Both proteins recognize the same core consensus sequence (TATCGATA).

In summary, CUT&RUN combined with csaw analysis identified 1,882 MCCS, of which 1,623 (86%) had not been previously characterized. These newly identified sites are predominantly promoter-proximal, in contrast to the intragenic localization of HAS and PionX sites. CUT&RUN generated sharp, focal MSL2 peaks compared to the broad peaks observed by ChIP. The native chromatin conditions of CUT&RUN, which omits crosslinking, indicates that many of these sites correspond to direct MSL2-DNA contacts, rather than indirect associations stabilized by crosslinking to neighboring bound factors.

### *roX2* is essential for chromosome binding of the DCC

Previous organismic studies suggested that in the absence of *roX* or the helicase maleless (MLE), the binding of MSL2 to the X chromosome is much reduced, but that a core complex of MSL2-MSL1 can bind to a small number of loci on the X chromosome (13,24,75). Furthermore, recombinant MSL2 is able to bind MRE motifs including ‘PionX’ signature sites *in vitro* in the absence of RNA (56,74). These observations suggested that *roX* RNA somehow modulates the intrinsic interactions of MSL2 to MREs *in vivo*. The availability of cell lines lacking *roX* RNA provides an opportunity to explore the *roX*-dependence of MSL2 binding.

We displayed the CUT&RUN signals for MSL2 and H4K16ac at 56 PionX sites, 309 HAS and all 1882 MCCS as separate heat maps, ordered according to their intensities in WT cells (Fig. 4A). The cumulative signals are displayed above the columns. In both *roX2* KO clones, MSL2 and H4K16ac signals were dramatically reduced at all site classes and this reduction was evident across the entire X chromosome in genome-browser views (Fig. 4B). The compromised MSL2 binding and H4K16ac signals in KO-B cells were restored to WT levels upon expression of FL *roX2*. Analysis of signal distributions across MSL2 target genes revealed the promoter-proximal enrichment of MCCS signals and the characteristic 3′ enrichment pattern of H4K16ac accumulation in WT cells, which was lost in knockout clones and restored upon rescue (Fig. 4C).

**Figure 4.**
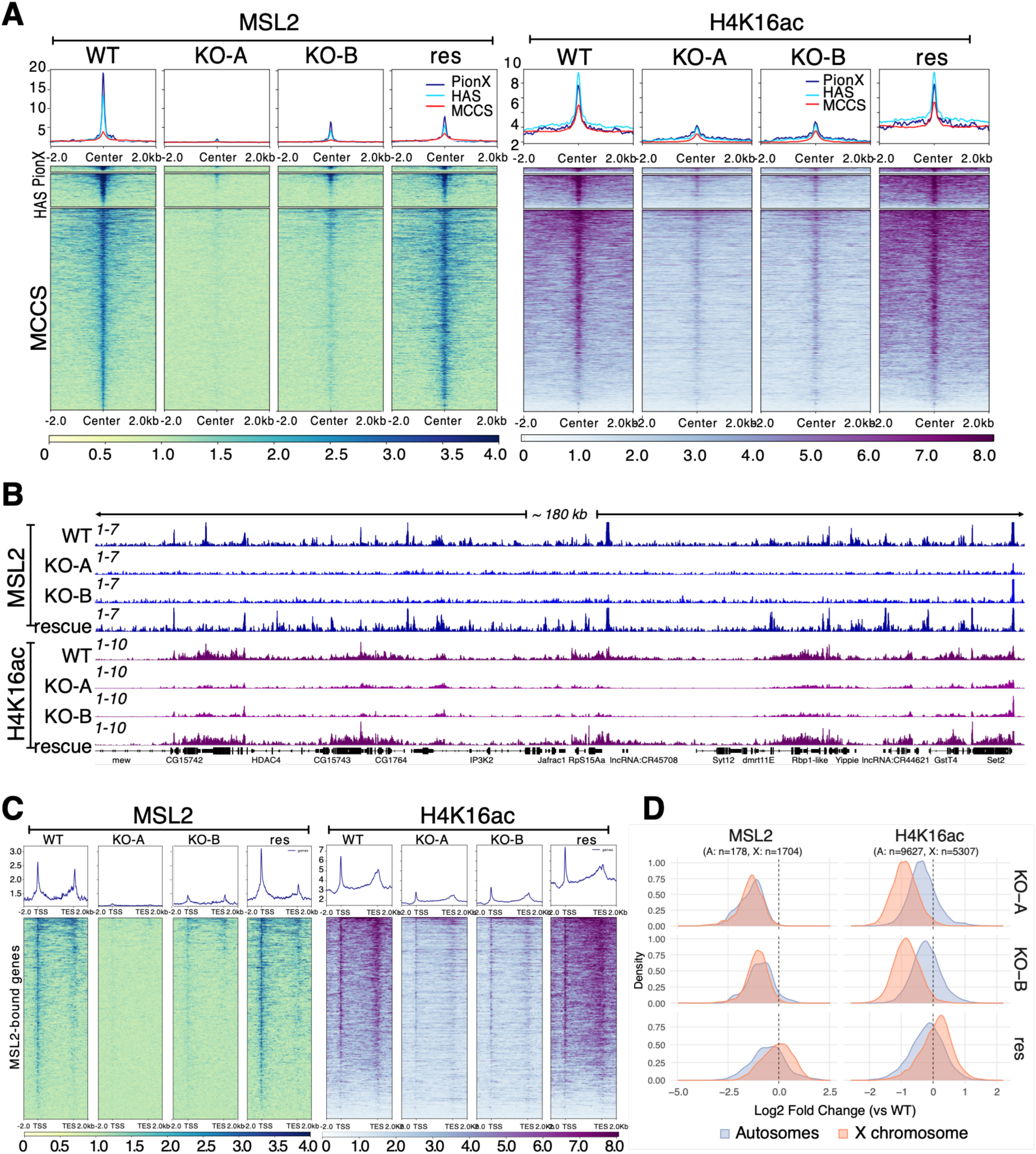
*Rox*-dependent chromosome binding of MSL2 (A) MSL2 and H4K16ac localization across site classes and genotypes. Heatmaps and metaplots depicting MSL2 (blue) and H4K16ac (purple) CUT&RUN signals normalized as ratio over negative control, centered on PionX (n=56, blue), HAS (n=309, cyan), and MCCS (n=1882, red) in WT, KO and rescue (res) cells. Sites are ordered by WT signal intensity. Signal is shown ±2 kb from peak centers. Scales are indicated at the bottom. (B) Representative genome browser coverage tracks for merged (n=3) MSL2 and H4K16ac CUT&RUN signals across the X-chromosome region of Fig. 3A in the indicated cell lines. (C) MSL2 and H4K16ac distribution at MSL2-bound genes. Heatmaps and metaplots showing MCCS and HCCS signals aligned to scaled genes classified as MSL2-bound. Gene bodies are scaled to equal length (5 kb) and flanked by 2 kb upstream and downstream regions. TSS and TES are indicated. Genes are ordered by decreasing MSL2 signal in the WT. Zero-signal regions were excluded and missing data were treated as zero. (D) Changes in X/autosome partitioning of MSL2 and H4K16ac signals. Density distributions of signal intensity differences (log2 fold-change relative to WT) for MSL2 and H4K16ac in KO-A, KO-B and rescue cells, as indicated. Differences were computed separately for autosomes (blue) and the X chromosome (red) at MCCS and HCCS sites, for MSL2 and H4K16ac respectively. The dashed line denotes no change. All signal intensities are mean CUT&RUN enrichments (target/control) extracted from merged biological replicates. In both knockout clones, X-chromosomal MSL2 and H4K16ac signals shift toward negative delta values, indicating reduced occupancy relative to WT, while autosomal signals remain centered near zero.

To further validate the global redistribution of the DCC upon *roX2* loss, we computed log2 fold-changes in MSL2 and H4K16ac signals relative to WT, separately for autosomal and X-chromosome sites (Fig. 4D). All chromosomal interactions of MSL2, whether on autosomes or the X chromosome, were diminished in both knockout clones. H4K16ac was selectively lost from the X chromosome, as expected from loss of dosage compensation, but autosomes did not gain corresponding signals. As before, *roX2* expression in the KO-B line fully restored X-chromosome enrichment, confirming that *roX2* is essential for proper genomic targeting of the DCC.

In summary, the analysis revealed a surprising requirement of *roX* RNA for all MSL2 chromosomal interactions in cells, independent of their sequence, shape, affinity or even chromosomal location. Evidently, the intrinsic DNA sequence readout of MSL2 does not suffice for stable chromosome interactions but requires additional input from non-coding RNA.

### Dosage compensation is maintained in *roX2* knockout cells

In view of the global loss of DCC binding to the X chromosome in *roX2* knockout cells, we wished to confirm that dosage compensation was impaired at the level of gene expression and identify *roX*-dependent gene expression changes. Towards this end, we established the transcriptomes of WT, KO and rescue cells through RNA-seq analyses. Principal component analysis (PCA) of all independent biological replicates confirmed the clear separation of the datasets according to genotype, and the absence of *roX2* in the KO clones (Supplementary Fig. 3A).

Despite the loss of *roX2,* all samples correlate highly with near perfect correlation for replicates of the same genotype (Supplementary Fig. 3B). Regardless, when comparing the KO and rescue conditions to the WT, most statistically significant gene expression changes differ according to cell line and not perturbation of the DCC per se (Supplementary Fig. 3C).

To our surprise, we found only minor global changes of transcription relative to WT, with log2 fold-change distributions for autosomal and X-linked genes centered around zero in KO-A, KO-B and rescue cells (Fig. 5A). Apparently, dosage compensation was largely maintained despite loss of a functional DCC from the X chromosome.

**Figure 5.**
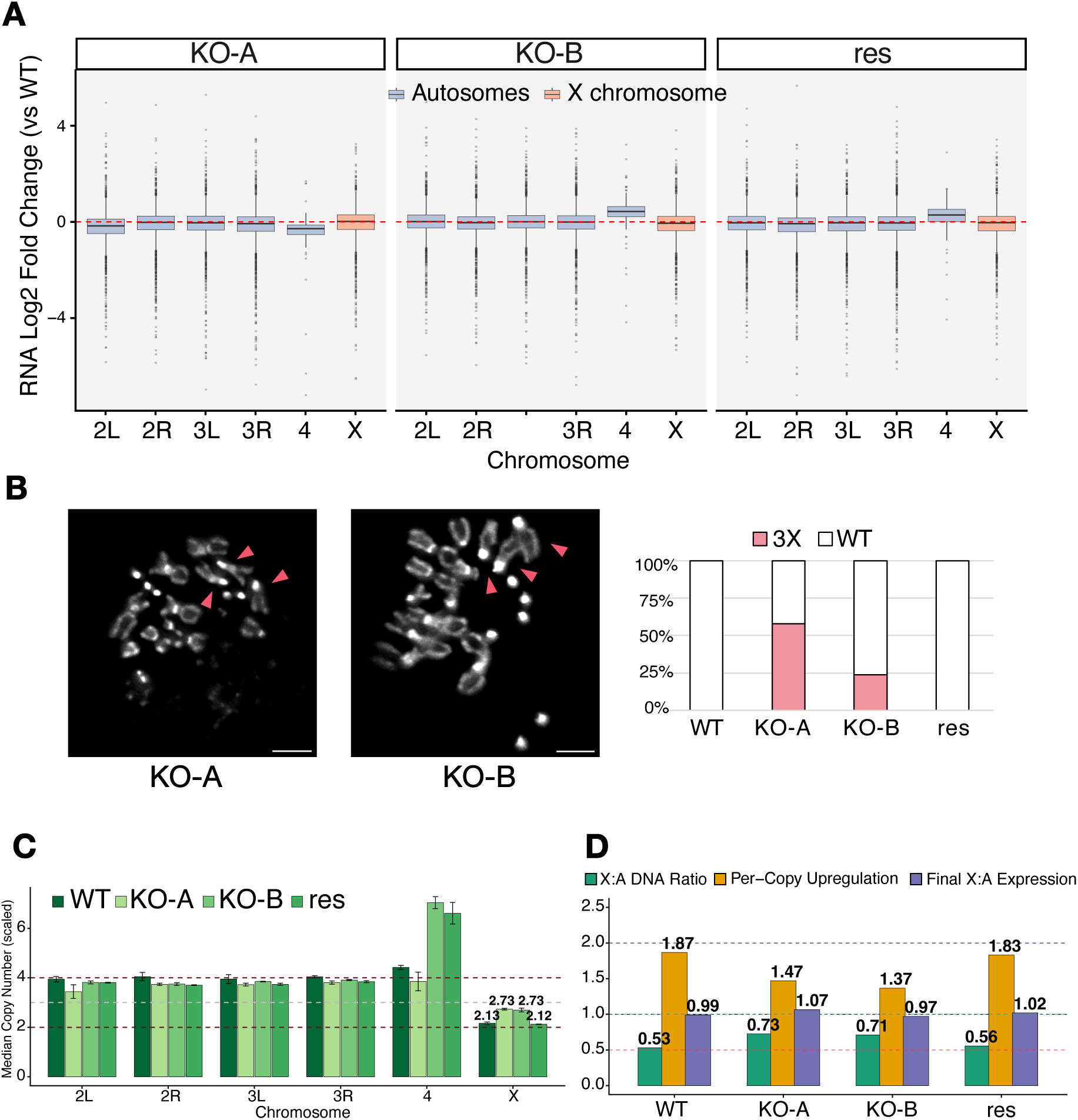
Dosage compensation is maintained in the roX2 KO cells. (A) RNA-seq analysis of gene expression changes for each chromosome. Boxplots show per-gene log2 fold-changes in RNA levels transcribed from protein-coding genes in KO-A, KO-B and rescue cells relative to WT, grouped by chromosomal arm (2L, 2R, 3L, 3R, 4, and X). Distributions for all chromosomes except the 4^th^ are centered near zero, indicating only very minor global expression changes. Boxes represent the interquartile range (IQR) with median lines; whiskers extend to 1.5 x IQR. The dashed red line marks no change (log2 fold change = 0). (B) Heterogeneity of karyotypes in KO cells. Left, exemplary images illustrating the existence of three X chromosomes (marked by pink arrows) in KO-A and KO-B. Scale bar, 5 μm. Right, barplot showing the percentage of cells having either 3 X chromosomes or the WT karyotype (see also Supplementary Table 2). (C) Chromosome copy number estimates derived from ChIP–MNase input sequencing. Median inferred copy numbers for the indicated cell lines are scaled to an autosomal baseline defined by the tetraploidy of WT cells (excluding the 4^th^ chromosome). Dashed lines denote the theoretical values. Both KO lines show elevated average X-chromosome copy number (values above the bars), whereas the rescue line restores X-chromosome copy number toward ∼2. Note the amplification of the 4^th^ chromosome in KO-B and its descendant rescue line. Error bars indicate SD across 3 independent biological replicates. (D) Dosage compensation by adjustment of X/autosome ratios. Bar plots show X/Autosome ratios: X:A DNA ratios from median inferred copy numbers as in C (green), X:A expression ratios corrected for DNA copy number (per-copy upregulation, yellow) and X:A expression ratios without correcting for DNA copy number (purple). The dashed lines indicate the theoretical ratios expected for full dosage compensation (X:A DNA 0.5, X:A copy number-corrected 2, X:A expression 1). The actual values are indicated above the bars.

To understand this unexpected finding, we performed a small-scale karyotype analysis of about 20 individual cells from each condition (see Supplementary Table 2 for a comprehensive summary of karyotypes). S2 cells are known to be male and mostly tetraploid, with a chromosome complement of four of each autosome and two X chromosomes (typical WT = 2xX, 4×2nd, 4×3rd, 2×4th and 1×2L = 13 chromosomes). We found the expected karyotype in all WT cells. Strikingly, about half of KO-A nuclei and about a quarter of KO-B nuclei contained a third X chromosome, which is clearly identified because of its acrocentric nature (representative images in Fig. 5B). We also found many cases, in which an autosome appeared to be missing, but these presumed aneuploidies were obscured by the presence of broken chromosomes in our karyotypes (Supplementary Table 2). The variability of karyotype between individual cells across all lines was best illustrated by the variable number of the 4^th^ chromosome, ranging from 2 to 7 copies. Remarkably, the wild-type karyotype with exactly two X chromosomes was recovered in all rescue cells (Fig. 5B). Despite the low sample numbers and some uncertainty about the identification of all autosomes, the results of this qualitative analysis provided an important clue to explain dosage compensation in the absence of DCC interactions with the X chromosome.

To substantiate these findings quantitatively, we determined the chromosome copy numbers at the population level by counting DNA reads in input samples of our ChIP analysis. Consistent with the single-cell karyotype data, both *roX2* knockout clones showed an average X-chromosome copy number of about 2.7 when scaled to the WT expected autosomal baseline, while the rescue line value was similar to WT (Fig. 5C). This increase in copy number of the X chromosome seems to be chromosome-wide rather than segmental (Supplementary Fig. 4A, B). This suggests that KO clones maintained balanced expression by altering the X/autosome copy number ratios upon impairment of DCC-mediated regulation of transcription (Fig. 5C). When expression ratios were corrected for DNA copy number (per-copy upregulation), the knockout clones showed reduced per-copy expression from the X chromosome, consistent with impaired DCC function. However, the overall X:A (X:Autosome) expression ratio remained near 1.0 (Fig. 5D), due to a combination of increases in X-chromosome sequences and the generic feedback mechanism described by Zhang and Oliver (28). The sequencing also confirmed the amplification of chromosome 4 in KO-B and its derived rescue line. Evidently, chromosome segregation in S2 cells is error-prone to an extent that allows reversible adjustment of chromosome content under selective pressure.

These findings demonstrate that dosage compensation remains essential even in aneuploid S2 cells under culture conditions. The recovery of cells with additional X chromosomes in both knockout clones, and the subsequent loss of the extra X upon *roX* restoration, strongly suggests that balanced genome expression provides a selective advantage during cell line establishment.

The error-prone chromosome segregation inherent to S2 cells, evidenced also by variable chromosome 4 copy numbers across all genotypes, apparently allows adjustment of chromosome content when cells experience prolonged selection pressure from impaired dosage compensation.

Together, these results establish *roX2* as essential for DCC function in cultured cells and demonstrate that the strong selective pressure to maintain dosage balance can drive rapid, reversible karyotypic changes, providing a sensitive experimental system for quantitative evaluation of *roX* RNA variants in chromosome targeting.

## Discussion

### Noncoding *roX* RNA is required for all chromosomal interactions of the DCC

Our CUT&RUN/csaw analysis identified over 1,800 MSL2 binding sites (MCCS), vastly expanding the catalog of DCC occupancy compared to previous studies that identified only a few hundred of the strongest binding sites (25,29,56). This improved detection is due to refinement of mapping by CUT&RUN and the implementation of csaw. CUT&RUN yields better signal-to-noise ratios, reduced background at active promoter regions and avoids fixation artefacts inherent to ChIP (76), with some degree of accessibility bias remaining (35,77,78). Compared to conventional peak callers, csaw’s window-based approach is more flexible and sensitive, allowing it to detect a broader range of enrichment events while delivering robust, statistical inference suitable for complex and replicated experimental designs (52,71).

The expanded MCCS catalog includes most of the previously characterized HAS and PionX sites, but in addition numerous MREs of presumed lower affinity. The enrichment of MCCS on the X chromosome demonstrates the robustness and specificity of our methodology in delineating the DCC binding landscape. Approximately half of these sites map within 1 kb of TSSs, while a significant fraction localize to introns and 3’UTRs of genes. This resonates with earlier lower-resolution ChIP studies that reported some of the HAS mapping to promoter-proximal regions, but a more substantial fraction at introns and 3’UTR (29,56). The gene-body binding on the X, which is mirrored by the H4K16ac distribution, is in line with a proposed mechanism of dosage compensation through enhanced processivity of transcription elongation (1). Tian et al. proposed that *roX* dynamically regulates DCC distribution by forming TES-TSS loops that facilitate local RNA Pol II access rather than long-range spreading (79). These observations are compatible with facilitated reinitiation of the RNA Pol II machinery that may contribute to the transcription boost.

The binding of DCC close to TSSs may reflect interactions of MSL2 with the CLAMP transcription factor, a ubiquitous cofactor of MSL2 that also recognizes MRE elements (25,36,56,74,79–81). Interestingly, sequence motif searches among the expanded set of MCCS identified not only the known MRE motif, but also found the motif (YTATCGATAR) significantly enriched at MCCS, which matches consensus binding sites for BEAF-32 and DREF. BEAF-32 (82–84) and DREF (85–87) are known to bind many promoters and insulators and act as boundary elements. Future studies are needed to distinguish whether these factors contribute directly to DCC recruitment or are confounding factors associated with accessible chromatin.

### Implications for X-chromosome targeting by the DCC

Our findings challenge ‘two-step’ models for DCC targeting on the X chromosome, which propose that initial binding occurs independently of *roX* RNA, followed by *roX*-dependent spreading over considerable distances (1). Key to these models is that binding sites differ qualitatively in that they are either *roX*-dependent or *roX*-independent. In contrast, the ‘affinity hierarchy’ model argues for a continuum of DCC binding sites with varying affinities. Spreading may then involve the local enrichment of DCC around sites of higher affinity, followed by the dynamic association with MRE elements or H3K36-methylated nucleosomes along the concentration gradient (12,23,24). It has been suggested that the dynamic association of *roX* RNA, mediated by the helicase MLE, may define cycles of chromatin release and ‘reloading’, which would facilitate the ‘spreading’ of the DCC to all potential binding sites in the neighborhood (18,88).

Our genome-wide mapping in *roX2* knockout S2 cells demonstrates that loss of *roX* RNA results in a dramatic reduction of MSL2 and H4K16ac occupancy not only at lower-affinity MREs, but also at HAS and PionX sites, which are known to be selectively bound by recombinant MSL2 in the absence of *roX* in vitro (56). Although H4K16ac enrichment from the X-chromosome is lost, the global levels of the mark remain largely unchanged because MOF maintains global H4K16ac levels on autosomes independently of *roX* RNA in the context of the non-specific lethal (NSL) complex (89,90).

In the absence of *roX*, MSL2 protein levels are moderately reduced, which may contribute to the observed global loss of binding. Under such limiting MSL2 conditions, one might predict preferential occupancy of the highest-affinity sites. Instead, we find that MSL2 binding is abrogated at all sites, including those previously classified as high-affinity. Evidently, *roX* RNA is required for MSL2 association at all genomic locations, independent of type and affinity, consistent with direct MSL2–*roX2* interactions that promote stable chromosome association (10,20,22). Accordingly, our data support a model in which the DCC, through MSL2-DNA recognition and cooperation with accessory factors such as CLAMP, engages its binding sites with a range of intrinsic affinities, and *roX* RNA stabilizes these interactions on the X chromosome. We observed earlier through ‘Fluorescence recovery after photobleaching’ (FRAP) studies that *roX2* endows the X chromosome-bound DCC with ‘low-diffusion’ properties that effectively trap the DCC on the X (10,21). In combination with the known homeostasis mechanisms that assure that the numbers of DCC do not exceed the binding capacity of sites on the X chromosome (1,15,69,91) such ‘trapping’ could be the basis of X chromosome-selective binding.

Taken together, our findings support models that pose that all chromosomal binding sites for the DCC require *roX* for robust interaction, independent of their DNA sequence signature or composition.

### A strategy for systematic structure-function analysis of *roX* RNA requirements for DCC loading

CRISPR-engineered *roX2* knockout S2 cells can form the basis of an experimental strategy to dissect the structural and functional elements that allow *roX* RNA to interact with MSL proteins and to target the complex to the X-chromosomal territory. S2 cells naturally depend on *roX2* for dosage compensation, allowing us to avoid the tedious generation of *roX1/roX2* double mutants that are not straight forward in the organism (14). Similar structure-function analyses in *Drosophila* commonly score ‘male viability’ as a readout for DCC functionality (11,17,18) and are complicated by developmental dynamics and potential cell-specific effects (92). We sought to test the cellular system by analyzing *roX*-derivaties which had previously been shown to support some male viability in corresponding rescue experiments in flies.

In our rescue experiments the robust DCC localization to the X chromosome, assessed by immunostaining and single-cell imaging analyses, was only observed with full-length *roX2* RNA. Despite overexpression, several truncated *roX2* variants as well as *roX1*-D3, constructs previously reported to support organismal viability (17,18), failed to establish faithful DCC targeting in our cellular system. This discrepancy and the karyotypic adaptation that occurred during the selection of *roX2*-deleted cells, highlight a tight dosage sensitivity of cultured cells that could not have been anticipated. It is possible that in flies even cryptic DCC function may be compatible with some viability due to compensatory buffering mechanisms, whereas the systematic image-based profiling employed here detects more nuanced deficiencies in DCC targeting.

The texture-based analysis of MSL protein staining in individual cells revealed subtle differences in nuclear staining for the different *roX* constructs, suggesting that the approach may be useful to classify intermediate phenotypes. The inability of certain *roX*-derivatives to support DCC targeting may reflect sequestration of MSL proteins to repetitive, heterochromatic regions (93,94) or other non-functional nuclear foci, patterns that might distinctly contribute to the classification texture-based image analysis.

### Dosage compensation is essential even in aneuploid cultured cells

Dosage compensation in *Drosophila* occurs throughout the organism, though its extent varies across tissues and cell types (92). Male S2 cells, an immortalized hematopoietic line, have served as a valuable model for studying dosage compensation mechanisms (28,29). These cells are predominantly tetraploid but exhibit substantial aneuploidy (28,44,95). While aneuploidy is typically detrimental at the organismal level due to cumulative gene dosage effects and regulatory network collapse (96,97), cultured cells display greater tolerance to chromosomal imbalances (28). This resilience initially led us to hypothesize that S2 cells might tolerate perturbation of the DCC. Indeed, transient RNAi-mediated depletion of MSL proteins does not impair S2 cell proliferation over experimental timescales of 6–9 days (27–29), raising the question of whether dosage compensation is truly essential in this context.

We therefore assumed that generating *roX*-deficient S2 cells would not pose any problem. However, generating a stable *roX2* gene deletion line, the first of its kind, turned out to be very challenging, in stark contrast to the ease of generating equivalent knockouts in female Kc cells (unpublished observations). The fact that the two *roX*-deficient lines we finally obtained during the lengthy selection process had increased X chromosome copy numbers demonstrates the critical requirement for maintaining a balanced X/autosome expression ratio even in aneuploid cultured cells. This is remarkable, given that the DCC compensates the 2-fold reduced X chromosome copy number by only a 1.4-fold boost in transcription (28). This dosage compensation is completely defective in the absence of *roX*, since MSL2 levels are diminished and DCC binding across all sites is dramatically lost.

### Mechanisms of karyotypic adaptation in *roX*-deficient cells

The gain of an additional X chromosome in *roX2* knockout cells represents a remarkable example of rapid evolution under selection. S2 cells are known to exhibit high rates of chromosome segregation defects during mitosis and variable chromosome copy numbers, particularly of chromosome 4 (28,44,95,98). Such defects arise from multiple mechanisms, including weakened spindle checkpoints and aberrant chromosome segregation (99). Like many tumor cells, S2 cells overexpress the protooncogene *ras* (95), which is known to trigger chromosome segregation defects and have multiple centromeres (100,101). Additionally, recent studies in mammalian cells have shown that disruption of the MSL complex and loss of H4K16ac primarily perturb the DNA replication program and promote chromosomal instability rather than directly altering transcription, highlighting the link between MSL/H4K16ac pathways and genome integrity (102).

Individual cells from Drosophila Genomics Resource Center (DGRC) lines, including S2 cells, display substantial karyotypic heterogeneity even within clonal populations (95). We assume that this was also true for the wild-type S2 cell population we used for our *roX* deletion experiments. During the extended selection process following CRISPR-Cas9 editing (∼6 weeks), cells that stochastically acquired a third X chromosome would have gained a selective advantage by restoring X/autosome expression balance without requiring a functional DCC. This hypothesis is strongly supported by our single-cell karyotyping and DNA copy number analyses, which showed an average of ∼2.7 X chromosomes in both independent knockout populations, suggesting this copy number represents an adaptive equilibrium that partially compensates for impaired DCC function while avoiding excessive aneuploidy-associated fitness costs. Heterogenous losses of individual autosomes likely also contribute to rebalancing the X/autosome expression ratio. We assume that, collectively, these karyotypic changes compensate for the loss of DCC-dependent dosage compensation.

The remaining compensation is likely achieved through a combination of gene-specific feedback mechanisms that operate whenever large parts of chromosomes are deleted (28,103,104) and post-transcriptional buffering mechanisms, including regulation of mRNA stability, translation and proteostasis (69,105,106). The gains of X chromosome in combination with such buffering mechanisms likely enabled knockout cells to stay competitive during selection despite complete loss of DCC function.

An important observation made through karyotyping of individual cells is that the established *roX*-depleted cells are no longer clonal but represent a mixed population of cells with either three X chromosomes or autosomal aneuploidies that arose from stochastic mitotic mis-segregation events in the wake of *roX2* gene deletion. This karyotypic heterogeneity also explains the reversibility of the phenotype upon re-expression of *roX2* in our rescue experiment, when the levels of MSL2 were partly restored and the binding of DCC to the X chromosomal territory re-established. Under these circumstances, an extra X chromosome is disadvantageous due to overcompensation and was selected against during the establishment of the rescue lines. This has been demonstrated in CHO cell lines, where cells under stringent phenotypic selection (e.g. expression of a transgene) are part of a more homogeneous population with fewer chromosome variations, while subcloning does not increase said homogeneity or chromosomal stability (107). The dynamics observed in cultured cells may not fully recapitulate organismal contexts. Nonetheless, the low-level viability of *roX*-null male flies (∼5% escapers *in vivo*;(13,14)) suggests that similar compensatory mechanisms might operate during development.

In summary, our findings establish *roX*-deficient S2 cells as a versatile and tractable platform for future structure–function studies of lncRNA-guided chromatin regulation in the context of dosage compensation. Reversible switching of dosage compensation provides a unique experimental opportunity for studying chromosomal evolution under controlled selective pressure. The unexpected karyotypic adaptation observed in these aneuploid cells upon *roX2* gene deletion underscores the existential importance of maintaining a balanced genome expression and the vital contribution of the DCC to this end. Our findings also reveal a surprising cellular plasticity in response to (epi)genetic perturbation and selection, which could be subject to further studies. Refinement of the experimental strategy by rapid and reversible switching of *roX2* expression may avoid adaptive effects that obscure the true functions of *roX* RNAs and will pave the way for high-resolution dissection of the elements and mechanisms by which *roX* RNA mediates DCC assembly, X-chromosome targeting, and epigenetic regulation.

## Supporting information

Gkountromichos_et_al_2026_Supplementary_data

## Acknowledgements

The authors gratefully acknowledge A. Thomae from the BMC Bioimaging Core Facility for training and invaluable advice on microscopy techniques and analyses. We thank D. Hörl from the Human Biology Division (LMU Biology) for early pilot classifications and analysis of microscopy images, T. Straub from the BMC Bioinformatics Unit for access to the computational cluster and advice on bioinformatics and helpful scripts and S. Krebs of LMU LAFUGA for Illumina sequencing. Also, we thank the IMB Microscopy and Histology CF for the image deconvolution in this work. Finally, we thank all current lab members for fruitful discussions. Schematics in the manuscript were created on BioRender.com.

## Author contributions

FG: Conceptualization, Investigation, Data curation, Formal analysis, Validation, Writing—original draft, Writing—review & editing; GY: Investigation, Formal analysis (Karyotype experiment), Writing—review & editing; MJ: Methodology (developed initial widefield microscopy analysis), Writing—review & editing; ACS: Investigation; MM: Writing—review & editing; PH: Supervision of GY, Formal analysis (Karyotype experiment), Writing—review & editing; PPB: Conceptualization, Funding acquisition, Supervision, Validation, Writing—original draft, Writing—review & editing;

## Data availability

Sequencing data generated in this study will be made publicly available prior to journal publication and accession numbers will be provided in a revised version of this preprint.

## References

1. Lucchesi, J.C. and Kuroda, M.I. (2015) Dosage compensation in Drosophila. Cold Spring Harb Perspect Biol, 7.

2. Makki, R. and Meller, V.H. (2021) When Down Is Up: Heterochromatin, Nuclear Organization and X Upregulation. Cells, 10.

3. Copur, O., Gorchakov, A., Finkl, K., Kuroda, M.I. and Muller, J. (2018) Sex-specific phenotypes of histone H4 point mutants establish dosage compensation as the critical function of H4K16 acetylation in Drosophila. Proc Natl Acad Sci U S A, 115, 13336–13341.

4. Franke, A. and Baker, B.S. (1999) The *roX1* and *roX2* RNAs Are Essential Components of the Compensasome, which Mediates Dosage Compensation in *Drosophila*. Molecular Cell, 4, 117–122.

5. Kelley, R.L., Meller, V.H., Gordadze, P.R., Roman, G., Davis, R.L. and Kuroda, M.I. (1999) Epigenetic Spreading of the Drosophila Dosage Compensation Complex from roX RNA Genes into Flanking Chromatin. Cell, 98, 513–522.

6. Meller, V.H. (2003) Initiation of dosage compensation in Drosophila embryos depends on expression of the roX RNAs. Mech Dev, 120, 759–767.

7. Deng, X. and Meller, V.H. (2006) roX RNAs are required for increased expression of X-linked genes in Drosophila melanogaster males. Genetics, 174, 1859–1866.

8. Meller, V.H., Wu, K.H., Roman, G., Kuroda, M.I. and Davis, R.L. (1997) roX1 RNA Paints the X Chromosome of Male Drosophila and Is Regulated by the Dosage Compensation System. Cell, 88, 445–457.

9. Meller, V.H., Joshi, S.S. and Deshpande, N. (2015) Modulation of Chromatin by Noncoding RNA. Annual Review of Genetics, 49, 673–695.

10. Villa, R., Jagtap, P.K.A., Thomae, A.W., Campos Sparr, A., Forne, I., Hennig, J., Straub, T. and Becker, P.B. (2021) Divergent evolution toward sex chromosome-specific gene regulation in Drosophila. Genes Dev, 35, 1055–1070.

11. Quinn, J.J., Zhang, Q.C., Georgiev, P., Ilik, I.A., Akhtar, A. and Chang, H.Y. (2016) Rapid evolutionary turnover underlies conserved lncRNA-genome interactions. Genes Dev, 30, 191–207.

12. Prayitno, K., Schauer, T., Regnard, C. and Becker, P.B. (2019) Progressive dosage compensation during Drosophila embryogenesis is reflected by gene arrangement. EMBO Rep, 20, e48138.

13. Meller, V.H. and Rattner, B.P. (2002) The roX genes encode redundant male-specific lethal transcripts required for targeting of the MSL complex. The EMBO Journal, 21, 1084–1091.

14. Kim, M., Faucillion, M.L. and Larsson, J. (2018) RNA-on-X 1 and 2 in Drosophila melanogaster fulfill separate functions in dosage compensation. PLoS Genet, 14, e1007842.

15. Oh, H., Park, Y. and Kuroda, M.I. (2003) Local spreading of MSL complexes from roX genes on the Drosophila X chromosome. Genes Dev, 17, 1334–1339.

16. Kuroda, M.I., Kernan, M.J., Kreber, R., Ganetzky, B. and Baker, B.S. (1991) The maleless protein associates with the X chromosome to regulate dosage compensation in drosophila. Cell, 66, 935–947.

17. Ilik, I.A., Quinn, J.J., Georgiev, P., Tavares-Cadete, F., Maticzka, D., Toscano, S., Wan, Y., Spitale, R.C., Luscombe, N., Backofen, R. et al. (2013) Tandem stem-loops in roX RNAs act together to mediate X chromosome dosage compensation in Drosophila. Mol Cell, 51, 156–173.

18. Ilik, I.A., Maticzka, D., Georgiev, P., Gutierrez, N.M., Backofen, R. and Akhtar, A. (2017) A mutually exclusive stem-loop arrangement in roX2 RNA is essential for X-chromosome regulation in Drosophila. Genes Dev, 31, 1973–1987.

19. Maenner, S., Muller, M., Frohlich, J., Langer, D. and Becker, P.B. (2013) ATP-dependent roX RNA remodeling by the helicase maleless enables specific association of MSL proteins. Mol Cell, 51, 174–184.

20. Muller, M., Schauer, T., Krause, S., Villa, R., Thomae, A.W. and Becker, P.B. (2020) Two-step mechanism for selective incorporation of lncRNA into a chromatin modifier. Nucleic Acids Res, 48, 7483–7501.

21. Straub, T., Neumann, M.F., Prestel, M., Kremmer, E., Kaether, C., Haass, C. and Becker, P.B. (2005) Stable chromosomal association of MSL2 defines a dosage-compensated nuclear compartment. Chromosoma, 114, 352–364.

22. Valsecchi, C.I.K., Basilicata, M.F., Georgiev, P., Gaub, A., Seyfferth, J., Kulkarni, T., Panhale, A., Semplicio, G., Manjunath, V., Holz, H. et al. (2021) RNA nucleation by MSL2 induces selective X chromosome compartmentalization. Nature, 589, 137–142.

23. Demakova, O.V., Kotlikova, I.V., Gordadze, P.R., Alekseyenko, A.A., Kuroda, M.I. and Zhimulev, I.F. (2003) The MSL complex levels are critical for its correct targeting to the chromosomes in Drosophila melanogaster. Chromosoma, 112, 103–115.

24. Dahlsveen, I.K., Gilfillan, G.D., Shelest, V.I., Lamm, R. and Becker, P.B. (2006) Targeting determinants of dosage compensation in Drosophila. PLoS Genet, 2, e5.

25. Eggers, N., Gkountromichos, F., Krause, S., Campos-Sparr, A. and Becker, P.B. (2023) Physical interaction between MSL2 and CLAMP assures direct cooperativity and prevents competition at composite binding sites. Nucleic Acids Res, 51, 9039–9054.

26. Baum, B. and Cherbas, L. (2008) In Dahmann, C. (ed.), Drosophila: Methods and Protocols. Humana Press, Totowa, NJ, pp. 391–424.

27. Hamada, F.N., Park, P.J., Gordadze, P.R. and Kuroda, M.I. (2005) Global regulation of X chromosomal genes by the MSL complex in Drosophila melanogaster. Genes Dev, 19, 2289–2294.

28. Zhang, Y., Malone, J.H., Powell, S.K., Periwal, V., Spana, E., Macalpine, D.M. and Oliver, B. (2010) Expression in aneuploid Drosophila S2 cells. PLoS Biol, 8, e1000320.

29. Straub, T., Zabel, A., Gilfillan, G.D., Feller, C. and Becker, P.B. (2013) Different chromatin interfaces of the Drosophila dosage compensation complex revealed by high-shear ChIP-seq. Genome Res, 23, 473–485.

30. Scacchetti, A., Schauer, T., Reim, A., Apostolou, Z., Campos Sparr, A., Krause, S., Heun, P., Wierer, M. and Becker, P.B. (2020) Drosophila SWR1 and NuA4 complexes are defined by DOMINO isoforms. Elife, 9.

31. Benchling. (2021–2025).

32. Doench, J.G., Fusi, N., Sullender, M., Hegde, M., Vaimberg, E.W., Donovan, K.F., Smith, I., Tothova, Z., Wilen, C., Orchard, R. et al. (2016) Optimized sgRNA design to maximize activity and minimize off-target effects of CRISPR-Cas9. Nat Biotechnol, 34, 184–191.

33. Gratz, S.J., Ukken, F.P., Rubinstein, C.D., Thiede, G., Donohue, L.K., Cummings, A.M. and O’Connor-Giles, K.M. (2014) Highly specific and efficient CRISPR/Cas9-catalyzed homology-directed repair in Drosophila. Genetics, 196, 961–971.

34. Team, R.C. (2025).

35. Meers, M.P., Bryson, T.D., Henikoff, J.G. and Henikoff, S. (2019) Improved CUT&RUN chromatin profiling tools. Elife, 8.

36. Albig, C., Tikhonova, E., Krause, S., Maksimenko, O., Regnard, C. and Becker, P.B. (2019) Factor cooperation for chromosome discrimination in Drosophila. Nucleic Acids Res, 47, 1706–1724.

37. Jayakrishnan, M., Havlova, M., Veverka, V., Regnard, C. and Becker, P.B. (2025) Genomic context-dependent histone H3K36 methylation by three Drosophila methyltransferases and implications for dedicated chromatin readers. Nucleic Acids Res, 53.

38. Liu, N., Hargreaves, V.V., Zhu, Q., Kurland, J.V., Hong, J., Kim, W., Sher, F., Macias-Trevino, C., Rogers, J.M., Kurita, R. et al. (2018) Direct Promoter Repression by BCL11A Controls the Fetal to Adult Hemoglobin Switch. Cell, 173, 430–442 e417.

39. Schindelin, J., Arganda-Carreras, I., Frise, E., Kaynig, V., Longair, M., Pietzsch, T., Preibisch, S., Rueden, C., Saalfeld, S., Schmid, B. et al. (2012) Fiji: an open-source platform for biological-image analysis. Nat Methods, 9, 676–682.

40. Pau, G., Fuchs, F., Sklyar, O., Boutros, M. and Huber, W. (2010) EBImage--an R package for image processing with applications to cellular phenotypes. Bioinformatics, 26, 979–981.

41. Wickham, H., Averick, M., Bryan, J., Chang, W., McGowan, L., François, R., Grolemund, G., Hayes, A., Henry, L., Hester, J. et al. (2019) Welcome to the Tidyverse. Journal of Open Source Software, 4.

42. McInnes, L., Healy, J., Saul, N. and Großberger, L. (2018) UMAP: Uniform Manifold Approximation and Projection. Journal of Open Source Software, 3.

43. Haralick, R.M., Shanmugam, K. and Dinstein, I. (1973) Textural Features for Image Classification. *IEEE Transactions on Systems, Man, and Cybernetics*, **SMC****-**3, 610–621.

44. Olszak, A.M., van Essen, D., Pereira, A.J., Diehl, S., Manke, T., Maiato, H., Saccani, S. and Heun, P. (2011) Heterochromatin boundaries are hotspots for de novo kinetochore formation. Nat Cell Biol, 13, 799–808.

45. Langmead, B. and Salzberg, S.L. (2012) Fast gapped-read alignment with Bowtie 2. Nat Methods, 9, 357–359.

46. Li, H., Handsaker, B., Wysoker, A., Fennell, T., Ruan, J., Homer, N., Marth, G., Abecasis, G., Durbin, R. and Genome Project Data Processing, S. (2009) The Sequence Alignment/Map format and SAMtools. Bioinformatics, 25, 2078–2079.

47. Quinlan, A.R. and Hall, I.M. (2010) BEDTools: a flexible suite of utilities for comparing genomic features. Bioinformatics, 26, 841–842.

48. Ramirez, F., Ryan, D.P., Gruning, B., Bhardwaj, V., Kilpert, F., Richter, A.S., Heyne, S., Dundar, F. and Manke, T. (2016) deepTools2: a next generation web server for deep-sequencing data analysis. Nucleic Acids Res, 44, W160–165.

49. Kent, W.J., Zweig, A.S., Barber, G., Hinrichs, A.S. and Karolchik, D. (2010) BigWig and BigBed: enabling browsing of large distributed datasets. Bioinformatics, 26, 2204–2207.

50. Zerbino, D.R., Johnson, N., Juettemann, T., Wilder, S.P. and Flicek, P. (2014) WiggleTools: parallel processing of large collections of genome-wide datasets for visualization and statistical analysis. Bioinformatics, 30, 1008–1009.

51. Thorvaldsdottir, H., Robinson, J.T. and Mesirov, J.P. (2013) Integrative Genomics Viewer (IGV): high-performance genomics data visualization and exploration. Brief Bioinform, 14, 178–192.

52. Lun, A.T. and Smyth, G.K. (2016) csaw: a Bioconductor package for differential binding analysis of ChIP-seq data using sliding windows. Nucleic Acids Res, 44, e45.

53. Robinson, M.D., McCarthy, D.J. and Smyth, G.K. (2010) edgeR: a Bioconductor package for differential expression analysis of digital gene expression data. Bioinformatics, 26, 139–140.

54. Lawrence, M., Huber, W., Pages, H., Aboyoun, P., Carlson, M., Gentleman, R., Morgan, M.T. and Carey, V.J. (2013) Software for computing and annotating genomic ranges. PLoS Comput Biol, 9, e1003118.

55. Yu, G., Wang, L.G. and He, Q.Y. (2015) ChIPseeker: an R/Bioconductor package for ChIP peak annotation, comparison and visualization. Bioinformatics, 31, 2382–2383.

56. Villa, R., Schauer, T., Smialowski, P., Straub, T. and Becker, P.B. (2016) PionX sites mark the X chromosome for dosage compensation. Nature, 537, 244–248.

57. Conway, J.R., Lex, A. and Gehlenborg, N. (2017) UpSetR: an R package for the visualization of intersecting sets and their properties. Bioinformatics, 33, 2938–2940.

58. Lawrence, M., Gentleman, R. and Carey, V. (2009) rtracklayer: an R package for interfacing with genome browsers. Bioinformatics, 25, 1841–1842.

59. Pagès, H.a.A. P. and Gentleman, R. and DebRoy, S. (2025), pp. R package version 2.76.70.

60. Bailey, T.L. (2021) STREME: accurate and versatile sequence motif discovery. Bioinformatics, 37, 2834–2840.

61. Tanaka, E., Bailey, T., Grant, C.E., Noble, W.S. and Keich, U. (2011) Improved similarity scores for comparing motifs. Bioinformatics, 27, 1603–1609.

62. Morgan, M.P., Hervé Obenchain, Valerie Hayden, Nathaniel. (2025). Bioconductor.

63. Bhardwaj, V., Heyne, S., Sikora, K., Rabbani, L., Rauer, M., Kilpert, F., Richter, A.S., Ryan, D.P. and Manke, T. (2019) snakePipes: facilitating flexible, scalable and integrative epigenomic analysis. Bioinformatics, 35, 4757–4759.

64. S., A. (2010).

65. Ewels, P., Magnusson, M., Lundin, S. and Kaller, M. (2016) MultiQC: summarize analysis results for multiple tools and samples in a single report. Bioinformatics, 32, 3047–3048.

66. Dobin, A., Davis, C.A., Schlesinger, F., Drenkow, J., Zaleski, C., Jha, S., Batut, P., Chaisson, M. and Gingeras, T.R. (2013) STAR: ultrafast universal RNA-seq aligner. Bioinformatics, 29, 15–21.

67. Love, M.I., Huber, W. and Anders, S. (2014) Moderated estimation of fold change and dispersion for RNA-seq data with DESeq2. Genome Biol, 15, 550.

68. Pedersen, T.L. (2025). R Package.

69. Villa, R., Forne, I., Muller, M., Imhof, A., Straub, T. and Becker, P.B. (2012) MSL2 combines sensor and effector functions in homeostatic control of the Drosophila dosage compensation machinery. Mol Cell, 48, 647–654.

70. Gilfillan, G.D., Straub, T., de Wit, E., Greil, F., Lamm, R., van Steensel, B. and Becker, P.B. (2006) Chromosome-wide gene-specific targeting of the Drosophila dosage compensation complex. Genes Dev, 20, 858–870.

71. Nooranikhojasteh, A., Tavallaee, G. and Orouji, E. (2025) Benchmarking peak calling methods for CUT&RUN. Bioinformatics, 41, btaf375.

72. Alekseyenko, A.A., Peng, S., Larschan, E., Gorchakov, A.A., Lee, O.K., Kharchenko, P., McGrath, S.D., Wang, C.I., Mardis, E.R., Park, P.J. et al. (2008) A sequence motif within chromatin entry sites directs MSL establishment on the Drosophila X chromosome. Cell, 134, 599–609.

73. Straub, T., Grimaud, C., Gilfillan, G.D., Mitterweger, A. and Becker, P.B. (2008) The chromosomal high-affinity binding sites for the Drosophila dosage compensation complex. PLoS Genet, 4, e1000302.

74. Eggers, N. and Becker, P.B. (2021) Cell-free genomics reveal intrinsic, cooperative and competitive determinants of chromatin interactions. Nucleic Acids Res, 49, 7602–7617.

75. Gu, W., Wei, X., Pannuti, A. and Lucchesi, J.C. (2000) Targeting the chromatin-remodeling MSL complex of Drosophila to its sites of action on the X chromosome requires both acetyl transferase and ATPase activities. The EMBO Journal, 19, 5202–5211.

76. Jain, D., Baldi, S., Zabel, A., Straub, T. and Becker, P.B. (2015) Active promoters give rise to false positive ‘Phantom Peaks’ in ChIP-seq experiments. Nucleic Acids Res, 43, 6959–6968.

77. Skene, P.J. and Henikoff, S. (2015) A simple method for generating high-resolution maps of genome-wide protein binding. Elife, 4, e09225.

78. Henikoff, S., Henikoff, J.G., Kaya-Okur, H.S. and Ahmad, K. (2020) Efficient chromatin accessibility mapping in situ by nucleosome-tethered tagmentation. Elife, 9.

79. Tian, S.Z., Yang, Y., Ning, D., Fang, K., Jing, K., Huang, G., Xu, Y., Yin, P., Huang, H., Chen, G. et al. (2024) 3D chromatin structures associated with ncRNA roX2 for hyperactivation and coactivation across the entire X chromosome. Science Advances, 10, eado5716.

80. Kuzu, G., Kaye, E.G., Chery, J., Siggers, T., Yang, L., Dobson, J.R., Boor, S., Bliss, J., Liu, W., Jogl, G. et al. (2016) Expansion of GA Dinucleotide Repeats Increases the Density of CLAMP Binding Sites on the X-Chromosome to Promote Drosophila Dosage Compensation. PLoS Genet, 12, e1006120.

81. Soruco, M.M., Chery, J., Bishop, E.P., Siggers, T., Tolstorukov, M.Y., Leydon, A.R., Sugden, A.U., Goebel, K., Feng, J., Xia, P. et al. (2013) The CLAMP protein links the MSL complex to the X chromosome during Drosophila dosage compensation. Genes Dev, 27, 1551–1556.

82. Jiang, N., Emberly, E., Cuvier, O. and Hart, C.M. (2009) Genome-Wide Mapping of Boundary Element-Associated Factor (BEAF) Binding Sites in Drosophila melanogaster Links BEAF to Transcription. Molecular and Cellular Biology, 29, 3556–3568.

83. Philip, P., Pettersson, F. and Stenberg, P. (2012) Sequence signatures involved in targeting the male-specific lethal complex to X-chromosomal genes in Drosophila melanogaster. BMC Genomics, 13, 97.

84. Negedu, S., Vidrine, E.J. and Hart, C.M. (2025) Characterization of core promoter activation by the Drosophila insulator-binding protein BEAF. Scientific Reports, 15, 37129.

85. Matsukage, A., Hirose, F., Yoo, M.-A. and Yamaguchi, M. (2008) The DRE/DREF transcriptional regulatory system: a master key for cell proliferation. Biochimica et Biophysica Acta (BBA) - Gene Regulatory Mechanisms, 1779, 81–89.

86. Heseding, C., Saumweber, H., Rathke, C. and Ehrenhofer-Murray, A.E. (2017) Widespread colocalization of the Drosophila histone acetyltransferase homolog MYST5 with DREF and insulator proteins at active genes. Chromosoma, 126, 165–178.

87. Tue, N.T., Yoshioka, Y., Mizoguchi, M., Yoshida, H., Zurita, M. and Yamaguchi, M. (2017) DREF plays multiple roles during Drosophila development. Biochimica et Biophysica Acta (BBA) - Gene Regulatory Mechanisms, 1860, 705–712.

88. Kiss, A.E., Venkatasubramani, A.V., Pathirana, D., Krause, S., Sparr, Aline C., Hasenauer, J., Imhof, A., Müller, M. and Becker, Peter B. (2024) Processivity and specificity of histone acetylation by the male-specific lethal complex. Nucleic Acids Research, 52, 4889–4905.

89. Feller, C., Prestel, M., Hartmann, H., Straub, T., Söding, J. and Becker, P.B. (2012) The MOF-containing NSL complex associates globally with housekeeping genes, but activates only a defined subset. Nucleic Acids Research, 40, 1509–1522.

90. Raja, S.J., Charapitsa, I., Conrad, T., Vaquerizas, J.M., Gebhardt, P., Holz, H., Kadlec, J., Fraterman, S., Luscombe, N.M. and Akhtar, A. (2010) The Nonspecific Lethal Complex Is a Transcriptional Regulator in Drosophila. Molecular Cell, 38, 827–841.

91. Park, Y., Kelley, R.L., Oh, H., Kuroda, M.I. and Meller, V.H. (2002) Extent of Chromatin Spreading Determined by roX RNA Recruitment of MSL Proteins. Science, 298, 1620–1623.

92. Pal, S., Oliver, B. and Przytycka, T.M. (2025) Cell-Type Specific Variation in X-Chromosome Dosage Compensation in Drosophila. microPublication Biology, 2025.

93. Li, F., Schiemann, A.H. and Scott, M.J. (2008) Incorporation of the Noncoding roX RNAs Alters the Chromatin-Binding Specificity of the Drosophila MSL1/MSL2 Complex. Molecular and Cellular Biology, 28, 1252–1264.

94. Figueiredo, M.L.A., Kim, M., Philip, P., Allgardsson, A., Stenberg, P. and Larsson, J. (2014) Non-coding roX RNAs Prevent the Binding of the MSL-complex to Heterochromatic Regions. PLOS Genetics, 10, e1004865.

95. Lee, H., McManus, C.J., Cho, D.-Y., Eaton, M., Renda, F., Somma, M.P., Cherbas, L., May, G., Powell, S., Zhang, D. et al. (2014) DNA copy number evolution in Drosophila cell lines. Genome Biology, 15, R70.

96. Cook, R.K., Christensen, S.J., Deal, J.A., Coburn, R.A., Deal, M.E., Gresens, J.M., Kaufman, T.C. and Cook, K.R. (2012) The generation of chromosomal deletions to provide extensive coverage and subdivision of the Drosophila melanogaster genome. Genome Biology, 13, R21.

97. Lee, H., Cho, D.Y., Whitworth, C., Eisman, R., Phelps, M., Roote, J., Kaufman, T., Cook, K., Russell, S., Przytycka, T. et al. (2016) Effects of Gene Dose, Chromatin, and Network Topology on Expression in Drosophila melanogaster. PLoS Genet, 12, e1006295.

98. Heun, P., Erhardt, S., Blower, M.D., Weiss, S., Skora, A.D. and Karpen, G.H. (2006) Mislocalization of the Drosophila centromere-specific histone CID promotes formation of functional ectopic kinetochores. Dev Cell, 10, 303–315.

99. Holland, A.J. and Cleveland, D.W. (2009) Boveri revisited: chromosomal instability, aneuploidy and tumorigenesis. Nat Rev Mol Cell Biol, 10, 478–487.

100. Chunduri, N.K. and Storchova, Z. (2019) The diverse consequences of aneuploidy. Nat Cell Biol, 21, 54–62.

101. Storchova, Z. and Kuffer, C. (2008) The consequences of tetraploidy and aneuploidy. Journal of Cell Science, 121, 3859–3866.

102. Milan, M., Runfola, V., Patel, M., Falbo, L., Noberini, R., Soriani, C., Rodighiero, S., Bonaldi, T., Costanzo, V., Pradeepa, M.M. et al. (2025) Mammalian H4K16ac regulates the spatiotemporal order of genome replication rather than gene expression. Nucleic Acids Res, 53.

103. McAnally, A.A. and Yampolsky, L.Y. (2009) Widespread transcriptional autosomal dosage compensation in Drosophila correlates with gene expression level. Genome Biol Evol, 2, 44–52.

104. Malone, J.H., Cho, D.-Y., Mattiuzzo, N.R., Artieri, C.G., Jiang, L., Dale, R.K., Smith, H.E., McDaniel, J., Munro, S., Salit, M. et al. (2012) Mediation of Drosophilaautosomal dosage effects and compensation by network interactions. Genome Biology, 13, R28.

105. Chen, Z.-X., Golovnina, K., Sultana, H., Kumar, S. and Oliver, B. (2014) Transcriptional effects of gene dose reduction. Biology of Sex Differences, 5, 5.

106. Zhang, Z. and Presgraves, D.C. (2017) Translational compensation of gene copy number alterations by aneuploidy in Drosophila melanogaster. Nucleic Acids Res, 45, 2986–2993.

107. Vcelar, S., Melcher, M., Auer, N., Hrdina, A., Puklowski, A., Leisch, F., Jadhav, V., Wenger, T., Baumann, M. and Borth, N. (2018) Changes in Chromosome Counts and Patterns in CHO Cell Lines upon Generation of Recombinant Cell Lines and Subcloning. Biotechnology Journal, 13, 1700495.

