## Supplementary material for "Dosage compensation defects due to *roX* RNA deletion are rescued by recalibration of X/autosome stoichiometry": Gkountromichos_et_al_2026_Supplementary_data

| Antibodies used in this study | | | | |
|---|---|---|---|---|
| Name | WB | IF | CUT&RUN/ChIP | Ref. |
| Rb α-MSL2 | 1:1000 | 1:400 | 1:500 | Fauth et. al. 2010, Eggers et. al. 2023 |
| Pre-immune serum (PPI) | | | 1:500 | Fauth et. al. 2010, Eggers et. al. 2023 |
| Rb α-H4K16ac | | 1:500 | 1:250 | MIllipore, 07-329 |
| Rb α-IgG | | | 1:250 | Cell Sign., 2729 |
| Rt α-MSL3 (1C9) | | 1:10 | | Morales et. al. 2004 |
| Gp α-MSL2 | 1:2000 | 1:500 | | Albig et. al. 2019 |
| Ms α-Lamin (T40) | 1:1000 | 1:100 | | Gift from H. Saumweber |
| Rb α-MOF | 1:1000 | | | Prestel et. al. 2010 |
| Rt α-MLE | 1:500 | | | Izzo et. al. 2007 |
| Ms α-H3 | 1:2000 | | | Abcam, ab10799 |
| IRDye α-guinea pig 680RD | 1:10000 | | | Licor Bio, 926-68077 |
| IRDye α-rabbit 680RD | 1:10000 | | | Licor Bio, 926-68071 |
| IRDye α-mouse 800CW | 1:10000 | | | Licor Bio, 926-32212 |

| Fluorophores used in this study | | | | |
|---|---|---|---|---|
| α-guinea pig Cy2 | | 1:200 | | Jackson Immuno, 106-225-003 |
| α-rat Cy3 | | 1:1000 | | Jackson Immuno, 712-165-153 |
| α-rabbit Alexa647 | | 1:500 | | Jackson Immuno, 711-605-152 |

| Oligos used in this study | |
|---|---|
| Name | Sequence (5' – 3') |
| sgRNA1 | GCTAGAGCAGCTAGATGTTG CGG |
| sgRNA3 | GGTGCTGGCTTAGAGAGAGATGG |
| sgRNA5 | AGAGCGAGATGACAATAGAGAGG |
| proX2_F2 | CGCTGGAGGCAAGGATATGAGC |
| rightCRISPRcheckrv | TCTGTTAATGCGCCACCCTT |
| proX2_seq_F | CAAGAGCAGTCGCAGCCTCGAG |
| qPCR_GAPDH_F | GGAGCCACCTATGACGAAAT |
| qPCR_GAPDH_R | GTAGCCCAGGATTCCCTTC |
| qPCR_roX2_25_F | GACGTGTAAAATGTTGCAAATTAAG |
| qPCR_roX2_25_R | TGACTGGTTAAGGCGCGTA |
| qPCR_miniroX_F | AACGTTCTCCGAAGCAAAAA |
| qPCR_miniroX_R | TGCGTTCCAAGACACATTTT |
| qPCR_roX1_d3_F | ATGAACACAGCCAAAGCAAG |
| qPCR_roX1_d3_R | GGCTCAGGCGTATAACGATT |

**Supplementary Table 1: List of reagents used in this study.** For antibodies, the table provides the name, host species (Rb: rabbit; Rt: rat; Gp: guinea pig; Ms: mouse), dilution used in Western blotting (WB), immunofluorescence microscopy (IF) or CUT&RUN/ ChIP. Sources or prior use are indicated under 'Ref'.

**WT**

| chrX | chr2 | chr3 | chr4 | chrExtra2L | Unknown/Broken |
|---|---|---|---|---|---|
| 2 | 4 | 4 | 4 | 1 | |
| 2 | 4 | 4 | 4 | 1 | |
| 2 | 3 | 4 | 3 | 1 | 2 |
| 2 | 3 | 4 | 4 | 1 | 2 |
| 2 | 3 | 4 | 4 | 1 | 2 |
| 2 | 3 | 4 | 5 | 1 | 2 |
| 2 | 4 | 4 | 3 | 1 | |
| 2 | 4 | 3 | 4 | 1 | 2 |
| 2 | 3 | 4 | 5 | 1 | 2 |
| 2 | 4 | 4 | 3 | 1 | |
| 2 | 4 | 4 | 3 | 1 | |
| 2 | 4 | 4 | 4 | 1 | |
| 2 | 3 | 4 | 4 | 1 | 2 |
| 2 | 3 | 4 | 6 | 1 | 2 |
| 2 | 3 | 4 | 2 | 1 | 2 |
| 2 | 4 | 4 | 3 | 1 | |
| 2 | 4 | 4 | 6 | 0 | |

**KO-A**

| chrX | chr2 | chr3 | chr4 | chrExtra2L | Unknown/Broken |
|---|---|---|---|---|---|
| 3 | 3 | 3 | 5 | 1 | |
| 2 | 4 | 4 | 5 | 1 | |
| 3 | 4 | 3 | 5 | 1 | |
| 2 | 3 | 4 | 5 | 1 | 2 |
| 3 | 3 | 4 | 4 | 1 | |
| 3 | 3 | 4 | 3 | 1 | |
| 3 | 3 | 4 | 4 | 2 | 1 |
| 2 | 3 | 4 | 5 | 1 | 2 |
| 2 | 3 | 4 | 7 | 1 | 3 |
| 2 | 4 | 4 | 2 | 1 | |
| 2 | 4 | 4 | 7 | 1 | |
| 3 | 3 | 3 | 5 | 1 | 2 |
| 3 | 2 | 4 | 6 | 1 | 2 |
| 3 | 3 | 4 | 3 | 1 | |
| 2 | 3 | 4 | 5 | 1 | 1 |
| 3 | 3 | 3 | 4 | 1 | 2 |
| 3 | 3 | 4 | 6 | 1 | 2 |
| 2 | 2 | 4 | 4 | 1 | 4 |
| 3 | 3 | 3 | 6 | 1 | 2 |

**KO-B**

| chrX | chr2 | chr3 | chr4 | chrExtra2L | Unknown/Broken |
|---|---|---|---|---|---|
| 3 | 3 | 4 | 6 | 1 | 2 |
| 2 | 4 | 4 | 4 | 1 | |
| 3 | 4 | 4 | 3 | 1 | |
| 2 | 4 | 4 | 6 | 1 | 1 |
| 2 | 3 | 4 | 5 | 1 | 2 |
| 3 | 3 | 4 | 1 | 1 | 2 |
| 2 | 3 | 4 | 2 | 1 | 2 |
| 2 | 3 | 3 | 4 | 1 | 1 |
| 2 | 4 | 4 | 4 | 1 | |
| 2 | 4 | 4 | 2 | 1 | |
| 2 | 4 | 3 | 5 | 1 | 1 |
| 2 | 4 | 4 | 3 | 1 | |
| 2 | 3 | 4 | 2 | 1 | 2 |
| 2 | 3 | 4 | 3 | 1 | 2 |
| 2 | 4 | 4 | 4 | 1 | |
| 3 | 3 | 3 | 5 | 0 | 2 |
| 2 | 4 | 4 | 7 | 1 | |
| 3 | 3 | 4 | 2 | 1 | |
| 2 | 4 | 4 | 5 | 1 | |
| 2 | 3 | 4 | 6 | 1 | 2 |
| 2 | 4 | 4 | 5 | 1 | |

**rescue**

| chrX | chr2 | chr3 | chr4 | chrExtra2L | Unknown/Broken |
|---|---|---|---|---|---|
| 2 | 3 | 4 | 4 | 1 | 2 |
| 2 | 4 | 3 | 3 | 1 | 2 |
| 2 | 4 | 4 | 6 | 1 | |
| 2 | 4 | 4 | 7 | 1 | |
| 2 | 3 | 4 | 2 | 0 | 2 |
| 2 | 4 | 4 | 3 | 1 | |
| 2 | 4 | 3 | 7 | 1 | 1 |
| 2 | 3 | 4 | 5 | 1 | 2 |
| 2 | 4 | 4 | 6 | 1 | |
| 2 | 3 | 3 | 7 | 1 | 2 |
| 2 | 3 | 4 | 4 | 1 | 2 |
| 2 | 4 | 3 | 8 | 1 | 2 |
| 2 | 4 | 4 | 4 | 1 | |
| 2 | 4 | 3 | 5 | 1 | 1 |
| 2 | 3 | 4 | 4 | 1 | 2 |
| 2 | 3 | 4 | 3 | 1 | 2 |
| 2 | 4 | 4 | 8 | 1 | |
| 2 | 3 | 4 | 4 | 1 | 2 |
| 2 | 4 | 4 | 7 | 1 | |
| 2 | 3 | 4 | 9 | 1 | 2 |

**Supplementary Table 2. Comprehensive karyotyping analysis of metaphase chromosome spreads across genotypes**. Each row represents an individual cell, and each column represents a specific chromosome or chromosomal abnormality. Chromosome copy numbers were counted for each chromosome (2, 3, 4 and X) across all genotypes (WT, KO-A, KO-B, and rescue). The "Extra 2L" column indicates additional copies of chromosome 2L observed in most cells. The "Unknown/Broken" column represents unidentified or fragmented chromosomal material that could not be reliably assigned to a specific chromosome, primarily due to technical limitations during spread preparation or imaging. Data are presented separately for each condition to assess chromosomal stability and ploidy changes upon *roX2* knockout and rescue.

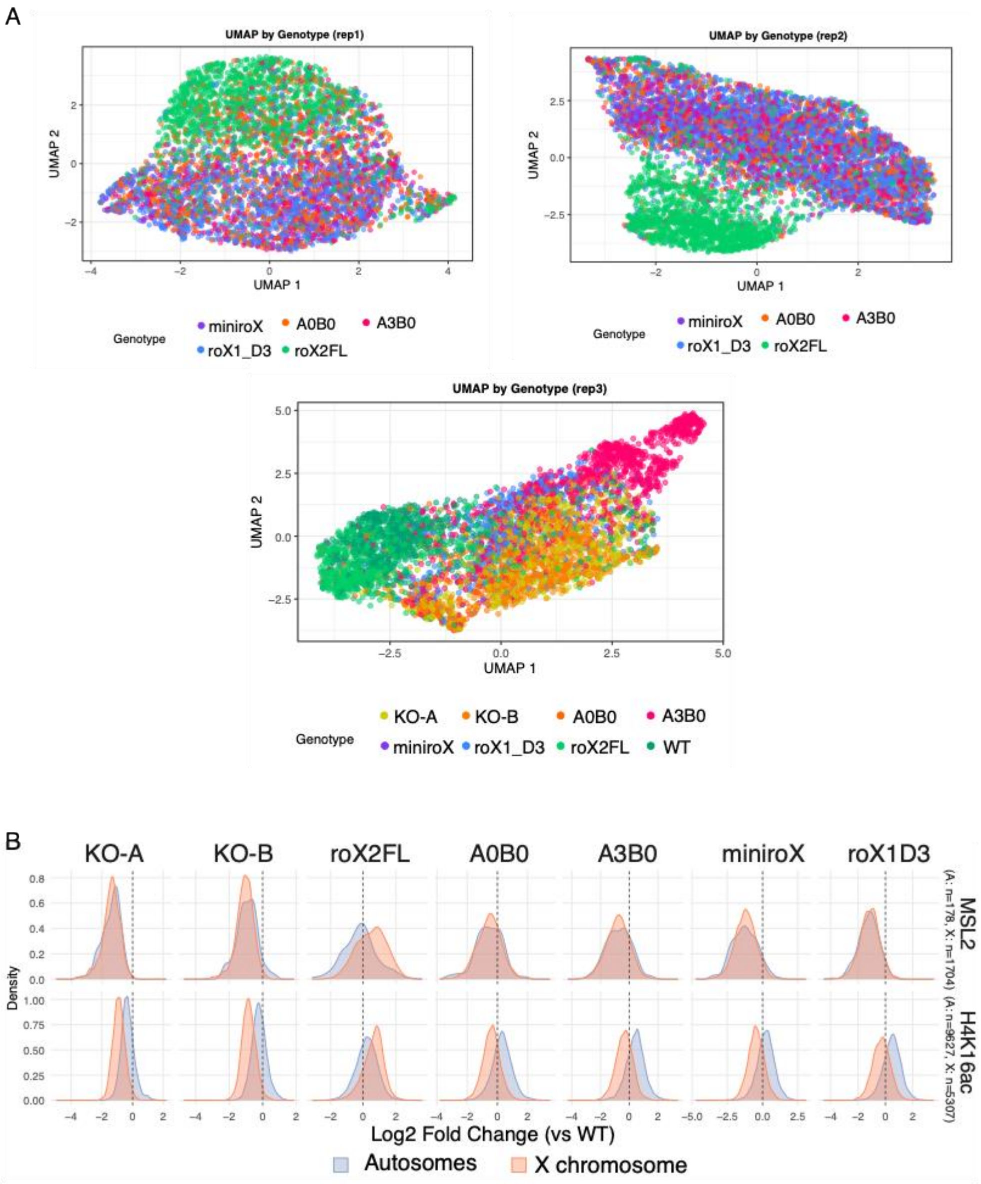


**Supplementary Figure 1: Replicate consistency of image-based MSL2 profiling and genomic assessment of rescue constructs**

(A) Classification of MSL2 localization states in single cells by image-based profiling. UMAP projection of single cells derived from widefield microscopy and quantified using MSL2 immunofluorescence intensity distributions and Haralick texture features. Each point represents one cell, colored by genotype/condition, as indicated. Three independent experiments [replicates (rep) 1-3] are shown. WT and *roX2FL* cell lines cluster together, indicative of the rescue capacity of the full-length construct.

(B) CUT&RUN quantification of X/autosome distribution changes of MSL2 (upper panel) and H4K16ac (lower panel) upon stable expression of *roX*-derivatives, as indicated. Changes in binding to autosomes (blue) and X chromosome (orange) (log2 fold-change relative to WT). The vertical dashed line indicates no change relative to WT. Constructs that partially rescue DCC targeting (roX2FL and *A0B0*) show some degree of X enrichment, whereas constructs (*A3B0*, *miniroX*, *roX1-D3*) do not support X chromosome targeting.

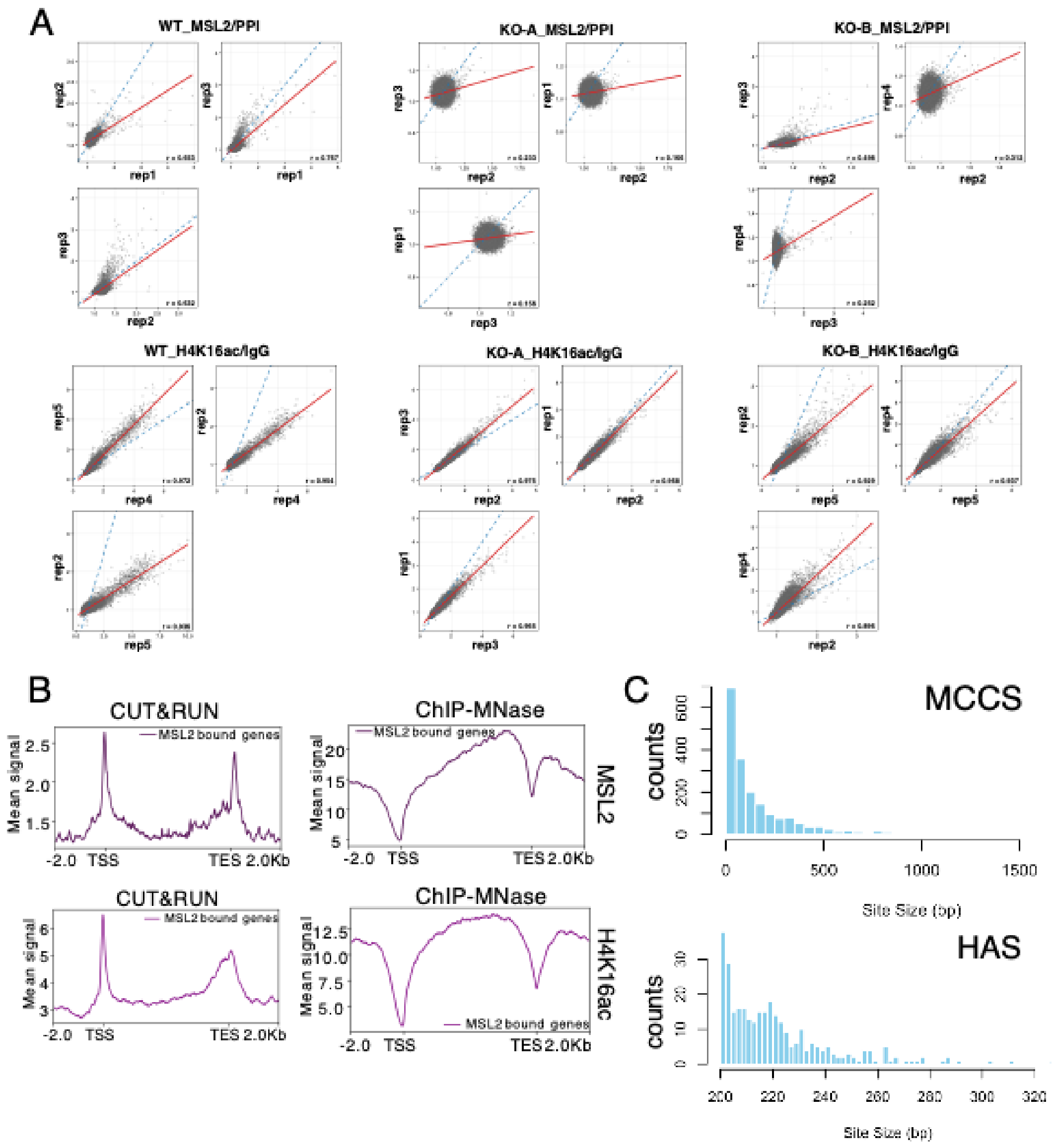


**Supplementary Figure 2: Biological replicate concordance and methodological comparison of CUT&RUN versus ChIP**

(A) Pairwise scatterplots showing genome-wide concordance between biological replicates of MSL2/PPI and H4K16ac/IgG ratio-normalized CUT&RUN data across wild-type (WT) and *roX2* knockout (KO-A and KO-B) S2 cells. Each point represents the normalized signal in one 10 kb genomic bin (dm6). Red line: linear regression; blue dashed line: perfect correlation (y = x). Pearson correlation coefficients (r) are shown in each panel. High correlations (H4K16ac: r > 0.92; MSL2 WT: r = 0.68) demonstrate excellent technical reproducibility. Lower MSL2 correlations in knockouts (r = 0.19-0.35) reflect biological variation from reduced protein abundance and unstable genomic binding, rather than technical noise.

(B) Method comparison at MSL2-bound genes. Meta-gene profiles comparing CUT&RUN (left) and ChIP (right) signal for MSL2 (top) and H4K16ac (bottom) in WT. CUT&RUN shows sharper peaks, lower background and lower affinity direct DNA binding, while ChIP reveals broader chromatin domains with higher signal. Genes bodies are scaled to 5 kb and ± 2 kb on either side of TSS and TES.

(C) Size distribution of the newly defined MSL2 CUT&RUN csaw sites (MCCS) and the canonical DCC binding sites, High-Affinity sites (HAS).

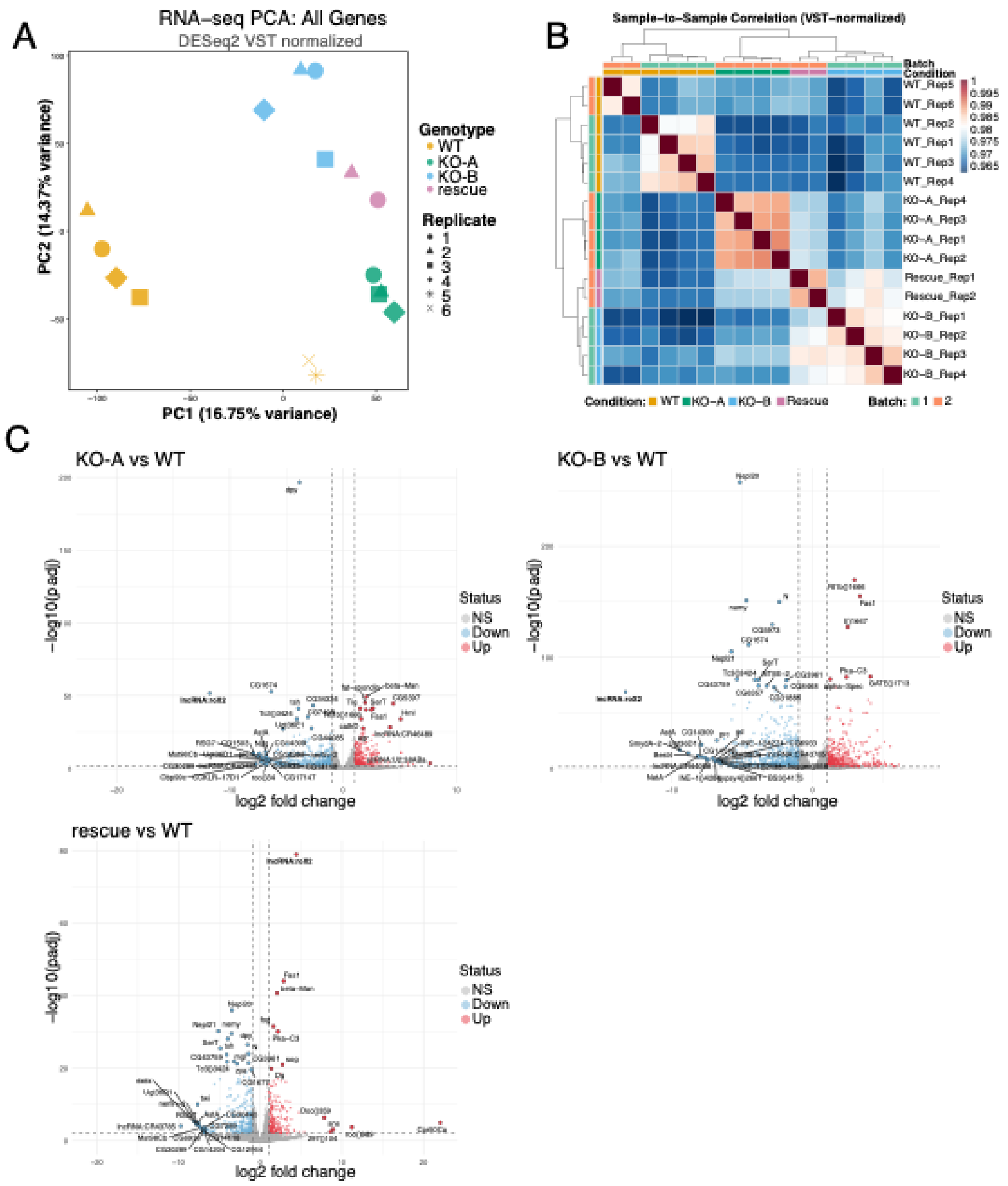


**Supplementary Figure 3: PCA, sample correlation, and differential expression analysis of *roX2* knockout transcriptomes**

(A) Principal component analysis (PCA) of VST-normalized RNA-seq data. Each shape represents one biological replicate; colors indicate genotype (WT = orange, KO-A = green, KO-B = sky blue, rescue = purple). PC1 (16.75%) separates samples by batch; PC2 (14.37%) captures genotype-specific differences.

(B) Sample correlation heatmap showing Pearson correlation coefficients between all RNA-seq samples based on VST-normalized expression (color: red = high, blue = lower). Hierarchical clustering uses Euclidean distance and Ward's linkage. All samples show high correlations (0.962–1.0, mean = 0.972), indicating excellent data quality. Clustering groups samples primarily by genotype rather than batch, demonstrating biological variation dominates over technical variation.

(C) Volcano plots of differential gene expression in pairwise comparison as indicated. Each point represents one gene (~20,000 genes). Vertical dashed lines mark $log_2FC| = 1$; horizontal line marks adjusted p-value = 0.01. Colors indicate differential expression status: upregulated (red), downregulated (blue), or not significant (gray). Top genes are labeled. Knockout lines exhibit widespread transcriptiosnal dysregulation affecting autosomes and X chromosome.

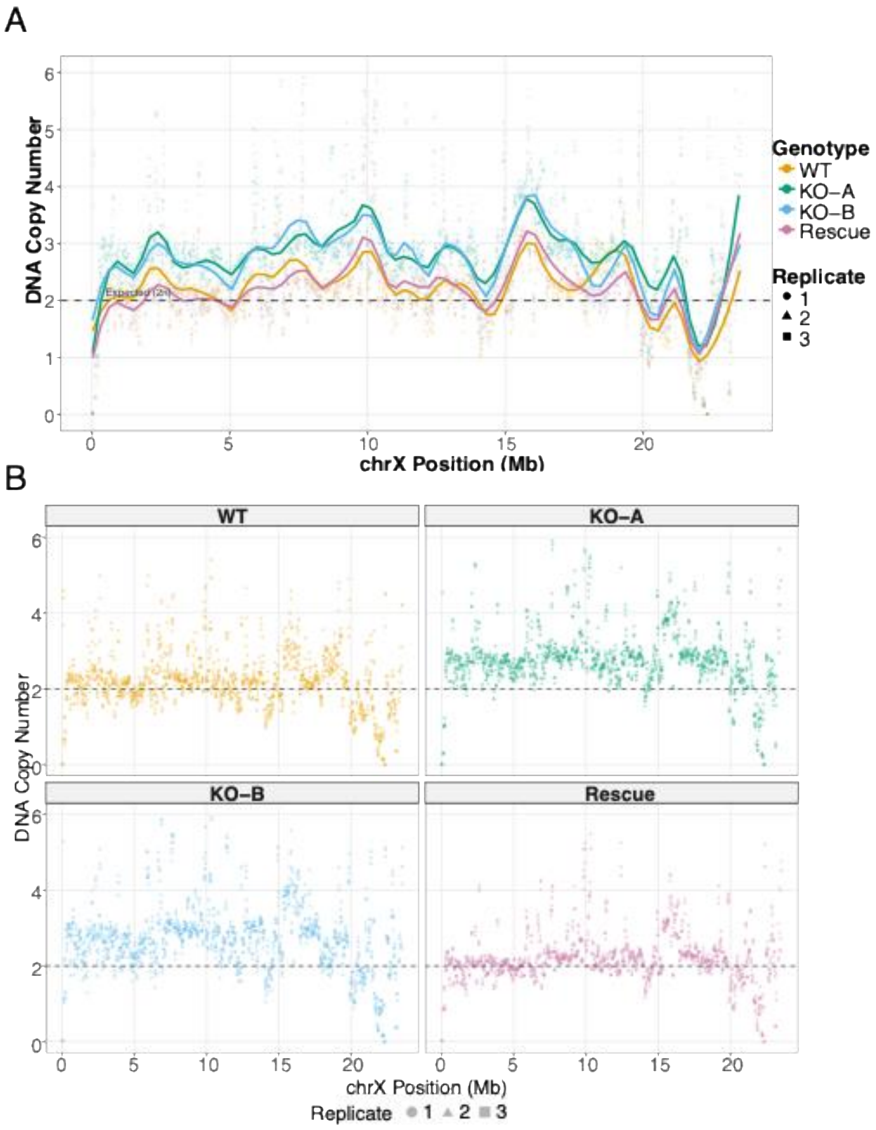


**Supplementary Figure 4: Copy number analysis demonstrates consistent X chromosome gains across independent *roX2* knockout clones**

(A) Smoothed chromosome X DNA copy number profiles across all conditions. Individual data points represent spike-in normalized copy number values for 50 kb genomic bins (n=471 bins per replicate). Genotypes are represented by different colors, replicates by shapes, as indicated. Smoothed trend lines (LOESS, span = 0.1) show the local average copy number for each genotype across the chromosome. The dashed horizontal line indicates the expected copy number for the X chromosome in male S2 cells (2n). Copy number values were scaled to absolute units using the wild-type autosomal baseline (see Methods). Both knockout clones (KO-A and KO-B) show consistently elevated X chromosome copy number (median ≈ 2.7–2.8 copies) compared to wild-type and rescue samples (median ≈ 2.2 copies), with the elevation distributed across the entire chromosome rather than localized to specific regions.

(B) Faceted view of chromosome X copy number by condition. The same bin-level data as in (A), displayed separately for each genotype to visualize replicate consistency. Points are colored by genotype and shaped by replicate. The dashed line indicates expected copy number (2n). Note the tight clustering of biological replicates within each condition and the consistent elevation in both independent knockout clones.
